# Marine protected areas promote resilience of kelp forests to marine heatwaves by preserving trophic cascades

**DOI:** 10.1101/2024.04.10.588833

**Authors:** Joy A. Kumagai, Maurice C. Goodman, Juan Carlos Villaseñor-Derbez, David S. Schoeman, Kyle C. Cavanuagh, Tom W. Bell, Fiorenza Micheli, Giulio De Leo, Nur Arafeh-Dalmau

**Author notes:** **Contact Information:** Joy A. Kumagai.

## Abstract

Under accelerating threats from climate change impacts, marine protected areas (MPAs) have been proposed as climate adaptation tools to enhance the resilience of marine ecosystems. Yet, debate persists as to whether and how MPAs may promote resilience to climate shocks. Here, we empirically assess whether a network of 85 temperate MPAs in coastal waters promotes resilience against marine heatwaves in Central and Southern California. We use 38 years of satellite-derived kelp cover to test whether MPAs enhance the resistance of kelp forest ecosystems to, and recovery from, the unprecedented 2014–2016 marine heatwave regime. We also leverage a 20-year time series of subtidal community surveys to understand whether protection and recovery of sea urchin predators within MPAs explain emergent patterns in kelp forest resilience through trophic cascades. We find that fully protected MPAs (i.e. no-take marine reserves) significantly enhance the resistance to and recovery of kelp forests to marine heatwaves in Southern California, but not in Central California. Differences in regional responses to the heatwaves may be partly explained by three-level trophic interactions comprising kelp, urchins, and predators of urchins. Urchin abundances in Southern California MPAs are significantly lower within fully protected MPAs during and after the heatwave, while the abundance of their predators are higher. In Central California, there is no significant difference in urchin abundances within protected areas as the current urchin predator, sea otters, are unilaterally protected. Therefore, we provide evidence that fully protected MPAs can be effective climate adaptation tools, but their ability to enhance resilience to extreme climate events depends upon region-specific environmental and ecological dynamics. As nations progress to protect 30% of the oceans by 2030 scientists and managers should consider whether protection will increase resilience to climate-change impacts given their local ecological contexts, and what additional measures may be needed.

## 1. Introduction

Marine protected areas (MPAs) are an essential conservation tool whose coverage has globally expanded in the past decades (Duarte et al., 2020; Lubchenco & Grorud-Colvert, 2015). Their importance is reflected in recent international policies aiming to protect 30% of coastal and open oceans, as specified within Target 3 of the post-2020 biodiversity framework (Convention of Biological Diversity, 2022). Following mounting evidence of increasing impacts of climate change on marine ecosystems (Schoeman et al., 2023), the new conservation framework includes climate mitigation and adaptation (e.g., Target 8 of the post-2020 biodiversity framework; Convention of Biological Diversity, 2022). The assumption underlying this framework is that protected areas may enhance climate adaptation and ecosystem resilience. While some empirical evidence supporting this expectation exists for individual MPAs and species (Jacquemont et al., 2022), clear empirical evidence at regional scales and for whole ecosystems is still lacking. There is strong consensus that well-managed and fully protected (i.e., no-take) MPAs promote biodiversity and habitat conservation (Gill et al., 2017; Lester et al., 2009; Sala & Giakoumi, 2018), but the extent to which MPAs confer ecological resilience to climate change impacts remains poorly understood.

One prominent manifestation of anthropogenic climate change is the increase in the frequency and intensity of extreme climate shocks, in particular marine heatwaves (MHWs) (Oliver et al., 2018). MHWs have caused mass mortality of sessile or low-mobility species (Garrabou et al., 2022; Szuwalski et al., 2023), losses of habitat-forming species such as corals and kelp, and regime shifts, among other impacts (Arafeh-Dalmau et al., 2019; McPherson et al., 2021; Smale et al., 2019; Wernberg, 2021). For example, MHWs in Australia and in the northeast Pacific Ocean have caused extensive losses of kelp over large areas and a shift into alternative stable ecosystem states dominated by less-productive algae or by sea urchin “barrens”, that have resulted in large-scale economic losses (Rogers-Bennett & Catton, 2019; Wernberg, 2021). Given that MHWs will become more frequent and intense in coming decades, it is a research priority to understand whether and how MPAs might increase resilience to these impacts.

Whether MPAs provide resilience to ecosystems experiencing climate shocks is debated and challenging to study. The operational definition for resilience used here is resistance to, and recovery from disturbance (Connell & Sousa, 1983), although resilience is a multifaceted concept (O’Leary et al., 2017). MPAs are designed to provide protection from local anthropogenic disturbance, primarily from extractive activities. They cannot directly mitigate the broad scale impacts of climate shocks, yet, by reducing extractive activities such as fishing, MPAs may allow the recovery of key species for ecosystem functioning, which in turn can promote resilience to climate shocks (Benedetti-Cecchi et al., 2024; Jacquemont et al., 2022; Roberts et al., 2017; Sala & Giakoumi, 2018; Schindler et al., 2015). The empirical evidence surrounding this argument is still emerging and mixed. Some studies have found no evidence that MPAs confer resilience to climate impacts (Bruno et al., 2018; Freedman et al., 2020; J. G. Smith et al., 2023). On the other hand, other studies have shown increased resilience to climate change in MPAs: for instance, in Baja California, Mexico, juvenile recruitment and adult abundance of pink and green abalone recovered faster within MPAs following a mass mortality of benthic invertebrates due to climate-driven hypoxia and warming (Micheli et al., 2012; A. Smith et al., 2022). In California, USA, species diversity recovered 75% faster from a series of MHWs within MPAs compared to adjacent unprotected areas (Ziegler et al., 2023). Additionally, a recent global analysis found that well-enforced MPAs can buffer the impacts of MHWs on reef fish by promoting the stability of fish at the community and metacommunity levels (Benedetti-Cecchi et al., 2024). Ultimately, a clear understanding of the conditions under which MPAs can provide climate resilience for whole ecosystems, including habitat-forming species and their associated communities, remains limited, due to the challenge of detecting resilience within MPAs.

One key challenge with detecting resilience emerges from the scarcity of long-term, sufficiently replicated and spatially extensive studies needed to characterize the state of the marine systems within and outside MPAs, before, during, and after climate extremes occur. Another limitation is that MPAs must be sufficiently large and must have been in place for a sufficient duration for any benefits of protection to emerge (Claudet et al., 2008). With a general paucity of studies with the necessary before, after, control, impact experimental design and statistical power, it is challenging to characterize the natural temporal variability and the inherent spatial heterogeneities of marine environments to achieve consensus on whether and under what circumstances MPAs might increase resilience to climate change impacts.

Here we overcome these challenges by utilizing long-term datasets to evaluate whether MPAs can increase kelp forest resilience to an unprecedented series of MHWs in California. During 2014–2016, the California coast was subject to one of the largest and longest MHW regime ever documented on Earth (Cavole et al., 2016; Di Lorenzo & Mantua, 2016; Frölicher & Laufkötter, 2018), providing a unique opportunity to investigate the dynamics of MPAs and ecosystem resilience. The combination of the 2014 warm-water anomaly and the 2015–2016 El Niño Southern Oscillation led to extremely warm waters (Cavole et al., 2016; Frölicher et al., 2018) that caused species range shifts (Favoretto et al., 2022; Sanford et al., 2019; J. G. Smith et al., 2023), a widespread loss of kelp forests from Northern California to Baja California Sur, Mexico (Bell et al., 2023), and an outbreak of sea urchins that are eroding kelp forest resilience. Additionally, California has a network of MPAs that cover 16% of state waters (Saarman & Carr, 2013), decades of satellite-derived estimates of kelp cover (Bell et al., 2023), and underwater surveys of kelp forest communities (Malone et al., 2022). With the rich ecological monitoring data that exist in this ecosystem, we can evaluate for the first time the resilience to and the underlying mechanisms of kelp forest ecosystems to MHWs within MPAs at a regional scale.

Trophic cascades are one of the proposed mechanisms by which MPAs can provide climate resilience. It has been hypothesized that, by protecting key predators of sea urchins, a voracious predator of kelp, MPAs may indirectly control sea urchin abundance, thus increasing both kelp resistance to, and recovery from, MHWs (Ripple et al., 2016). Outside MPAs, where fishers target urchin predators, including California sheephead (*Semicossyphus pulcher*) and spiny lobsters (*Panulirus interruptus*), there are fewer urchin predators and more urchins (Eisaguirre et al., 2020). When a disturbance leads to severe kelp loss, urchins may shift their behavior from hiding in protective cracks and eating drift kelp to being more exposed, eating any remaining kelp and preventing further kelp establishment (Harrold & Reed, 1985; Kriegisch et al., 2019). Overharvesting and depletion of urchin predators can then lead to a high abundance of urchins that overgraze kelp forests (Cowen, 1983). If MPAs protect and foster greater abundances of urchin predators (which otherwise would be commonly fished), then protected kelp forests may be more likely to recover and even resist change in the face of a disturbance, compared to unprotected kelp forests.

In this study we investigated the recovery of the giant kelp, *Macrocystis pyrifera* (henceforth “kelp”) following the 2014–2016 MHWs in Central and Southern California. The main objectives were to determine (1) whether kelp forests within a network of MPAs were more resilient to the 2014–2016 MHWs compared to unprotected kelp forests, (2) whether resilience of kelp forests differed between regions, and (3) whether there is evidence that trophic cascades are a mechanism underlying resilience to climate shocks. To address these questions, we assessed changes in kelp area during and after the 2014– 2016 MHW using satellite-derived estimates of kelp area spanning 1984–2021 and analyzed 20 years of subtidal monitoring datasets to investigate possible evidence for trophic cascades. We tested the following hypotheses: (i) kelp canopy resilience is higher within fully protected and partially protected areas compared to unprotected areas in both Central and Southern California during and after the MHWs; (ii) urchin abundances are lower within protected areas compared to unprotected areas during and after the MHWs, enabling the recovery of kelp forests; and (iii) urchin abundances are driven by the abundances of their main predators.

## 2. Materials and Methods

### 2.1 Study area

Our study spans Central and Southern California as defined by the Marine Life Protection Act (Marine Life Protection Act, 2013), encompassing the region where giant kelp is the dominant surface canopy-forming kelp species in the USA, from the US Mexico border (∼32.5° N) to Pigeon Point, California (∼37.2° N) (Figure 1). Central and Southern California are separated into two different biogeographic regions at Point Conception (∼34.5° N), which is a transition zone between the cooler temperate ecosystems of Central California and the warmer ecosystems of Southern California (Murray & Abbott, 1980). In this region, there are a total of 85 MPAs, with varying levels of fishing restriction (no-take MPAs and partially protected multiple-use areas) (Figure 1). Of these, 60 MPAs are in Southern California and 25 in Central California. In Southern California, the primary predators of sea urchins include the California sheephead and spiny lobsters, while in Central California, sea otters (*Enhydra lutris*) and sunflower sea stars (*Pycnopodia helianthoides*) are the primary predators of urchins — although sunflower sea stars are no longer a functional predator of urchins due to a mass mortality event that has greatly reduced their numbers throughout California (Burt et al., 2018; Eisaguirre et al., 2020). As such, sea otters, which are protected statewide, are currently the sole top predator of urchins within Central California and are known to be a primary driver for changes in kelp (Eisaguirre et al., 2020; Nicholson et al., 2024). California sheephead and spiny lobsters, which are both fished, fill this role of predation in Southern California (Eisaguirre et al., 2020). The range of purple (*Strongylocentrotus purpuratus*) and red urchins (*Mesocentrotus franciscanus*) both span from Alaska to Baja California, while crowned urchins (*Centrostephanus coronatus*) are common between the Galapagos and the Channel Islands, being mostly absent from Central California.

**Figure 1:**
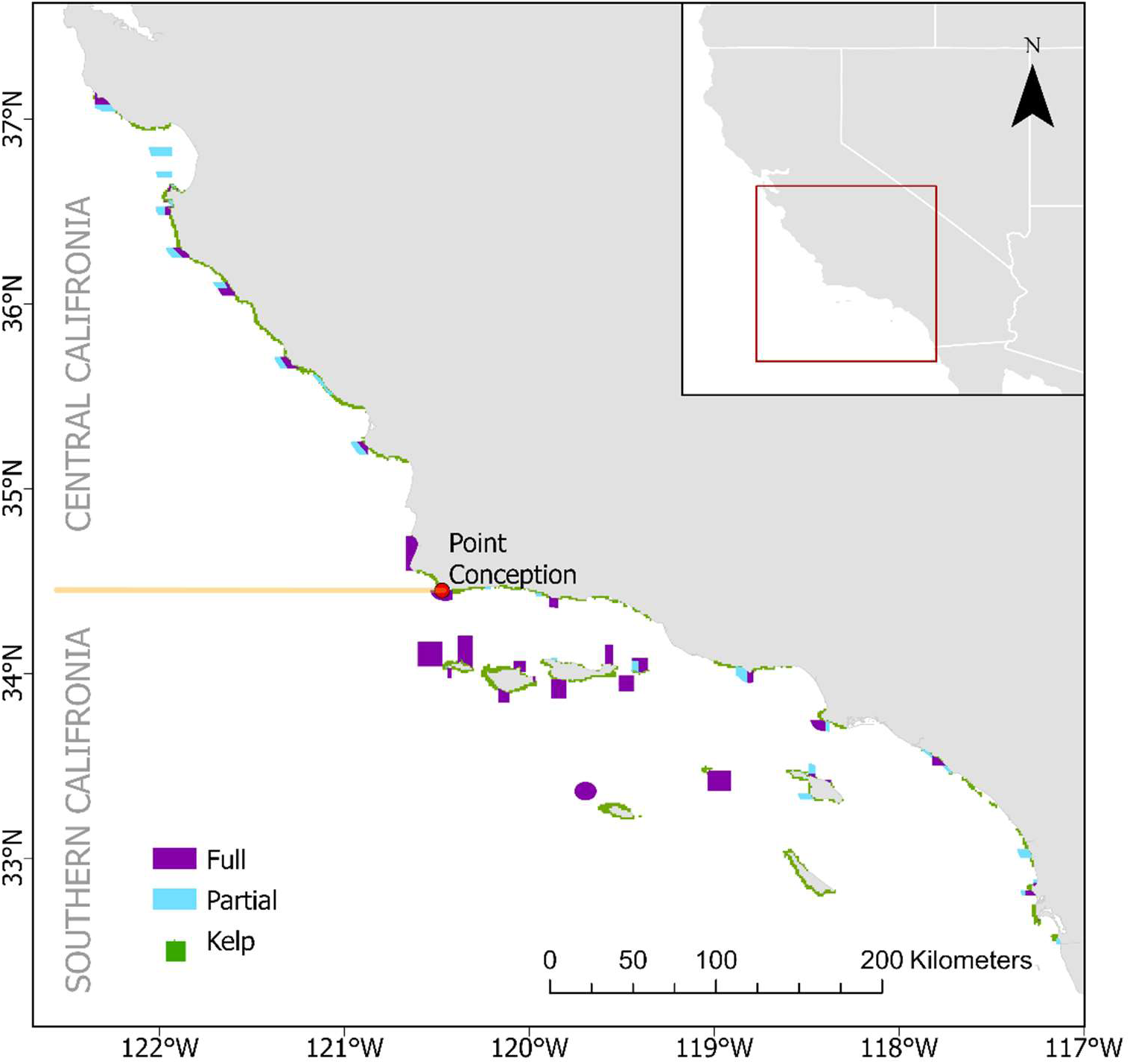
Distribution of giant kelp and the network of marine protected areas in Central and Southern California. Green polygons show the satellite-derived locations with giant kelp, and purple and blue polygons show fully and partially protected areas in the network in Central and Southern California. The yellow horizontal line at 34.4° N represents the biogeographic barrier at Point Conception where Central California is separated from Southern California.

### 2.2 Quantifying the resilience of kelp forests from MHWs

We used 38 years (1984–2021) of quarterly estimates of kelp area based on remote sensing from the Santa Barbara Coastal LTER time series dataset (Bell et al. 2023) to estimate the resilience of kelp forests. The dataset contains quarterly estimates of kelp canopy area in m^2^ (referred to as kelp area from now on) from three Landsat sensors: Landsat 5 Thematic Mapper (1984–2011), Landsat 7 Enhanced Thematic Mapper+ (1999-present), and Landsat 8 Operational Land Imager (2013-present). Each Landsat sensor has 30-m resolution and does not distinguish between giant and bull kelp. We aggregated the original dataset to 1-km resolution to reduce spatial autocorrelation in the data by summing the kelp area in the 30×30m pixels. We followed previous approaches for cleaning the Landsat data (Bell et al., 2023) and excluded those quarters of a year that had no data for more than 25% of the 30-m pixels. We also removed from our dataset 1-km pixels which consisted of fewer than five 30-m pixels. Next, we removed 1-km pixels for which more than two quarters of kelp area were missing in a given year. Finally, the quarterly 1-km data were aggregated to a yearly scale by taking the maximum quarterly kelp area for each year, as a preliminary data analysis showed that the peak in kelp forest cover might occur in different quarters in different years and pixels. Our final dataset thus uses the maximum annual kelp area per 1-km^2^ pixel.

To develop a metric of kelp resistance and recovery to the 2014–2016 MHWs, we calculated the relative change in kelp area. For each 1-km^2^ pixel, we first determined the long-term historic baseline of kelp coverage, defined as the average kelp area across the 30 years (1984–2013) before the 2014-16 MHWs. Next, we calculated the ratio of each subsequent year’s (2014–2021) kelp area relative to that baseline. We define the resistance and recovery of kelp forests to the MHWs as the annual change in kelp area during 2014–2016 and 2017–2021, respectively, relative to the corresponding pre-2014, historical baseline mean for each 1-km^2^ pixel. Accordingly, values close to 100% represented stable kelp cover with respect to the average kelp forest cover during the 1984–2013 baseline; values <100% represented kelp decline with respect to the pre-MHW baseline, and values >100% represented expansion of kelp coverage with respect to the historical baseline.

### 2.3 Evaluating the resilience of kelp forests within MPAs

#### 2.3.1 The MPA dataset

We downloaded the spatial layers, age, and level of fishing restriction for California’s MPAs from NOAA’s MPAs Inventory. We categorized MPAs as fully protected or partially protected. Fully protected areas do not allow any extractive activities, while there are some restrictions on recreational and commercial fishing within partially protected areas (multiple use areas). Next, we overlaid the MPA layer on the 1-km^2^ resolution kelp layer. This procedure allowed us to categorize the level of protection of each pixel as (i) unprotected, (ii) partially protected, or (iii) fully protected. Any pixels within National Marine Sanctuaries were classified as unprotected because many of these sanctuaries have minimal or no fishing restrictions. We then classified the remaining 85 MPAs in two age categories based on their year of implementation. We classified MPAs established before 2007 as “old” and those established between 2007 and 2012 as “new”. We also estimated additional environmental and human impact variables to investigate whether MPAs were established in more productive areas for kelp forests using a principal component analysis (Supplementary methods and results).

#### 2.3.2. Permutation Analysis

We used a one-tailed permutation analysis to test whether the differences in resistance and recovery of kelp area during and after the 2014–2016 MHWs were affected by protection status, i.e., fully protected vs partially protected vs unprotected areas. As there are known latitudinal differences in water temperature, oceanographic regimes and in other social and environmental drivers of kelp coverage, we repeated the analysis for Southern and Central California separately. Given the high year-to-year variability in kelp cover, we used a permutation test because it does not make assumptions about the underlying distribution of the data (Supplementary Figure 2). Specifically, we tested the following hypotheses: (i) relative kelp area during and after the MHWs within fully protected areas is higher than relative kelp area within partially protected or unprotected areas, and (ii) relative kelp area during and after the MHWs within partially protected areas is higher than that in unprotected areas.

For each region, we first computed the observed differences in the medians of the relative kelp area during the response period (2014–2016) and in the recovery period (2017–2021) for each category (i.e., fully protected vs unprotected; partially protected vs unprotected, fully protected vs partially protected). Next, to derive the null distribution, we randomly assigned each pixel to one of the three protection categories and computed the differences in the median values among the three categories of the randomized set as above. These values were saved and then the same calculation was replicated 10,000 times, each time randomly assigning each pixel a protection category. The respective null distributions of the difference in the median values among the three categories were derived by using the 10,000 randomized replicates, and a one-sided pseudo *p*-value was calculated as 1 less than the percentile of the observed value under the corresponding null distribution. Since we generated multiple *p*-values for each hypothesis, we applied Bonferroni’s correction, multiplying *p*-values by the number of comparisons undertaken (six). This analysis was implemented first across the entire study area, then repeated for each region individually. We also explored the effect of the age of MPAs on our results, repeating the permutation analyses for both old (established before 2007) and new (established between 2007-2012) MPAs separately.

The distribution of relative kelp area was highly right skewed with most pixels having kelp coverage after the MHW equal to, or lower than, before MHW. However, in some pixels, the relative differences in the median coverage during and after MHW with respect to the historical baseline exceeded 100% by several orders of magnitude. These substantial changes in kelp area reflect the fact that some areas that contained very little kelp historically experienced a large increase in kelp area during 2014– 2021. To test the impact of pixels with very small pre-MHW kelp forest area on the results of the permutation analysis, we conducted a sensitivity analysis that involved removing pixels with the lowest 5% to 30% of mean historic kelp area from the analysis in increments of 5% (Supplementary Table 6) and then re-running the permutation analysis.

### 2.4 Mechanism of resilience: trophic cascades

#### 2.4.1 Processing of subtidal dataset

To investigate whether species interactions - sea urchin grazing and trophic cascades - may be a mechanism driving differences in kelp recovery between protected and unprotected areas, we used subtidal surveys of kelp forest communities that include urchins and their main predators from the Partnership for Interdisciplinary Studies of Coastal Oceans (PISCO; Malone et al., 2022). We spatially joined the master PISCO sites dataset within our study area with the MPA layer to produce a layer with the sites, protection status, and region. Next, we created a dataset of all the unique transects where PISCO divers surveyed our species of interest for both the fish and benthic invertebrate (swath) surveys. We filtered the PISCO data from 2002–2023. We chose this start year because the UCSB and UCSC (University of California Santa Barbara and Santa Cruz) monitoring teams started consistently searching for crowned urchins using the same methods in 2002 and this year is also five years after the 1997–1998 extreme El Niño. We chose five years after the 97-98 period to ensure any effects of sheephead recruitment from the El Niño had mostly dissipated. We terminated the series in 2023 because this was the last year of available data. Additionally, we focused on adult organisms, and did not include urchin recruits and California sheephead that were <10 cm in total length, as these are not physically big enough to eat large urchins. A previous study found only very small sea urchin spines in the gut contents of California sheephead 15-20 cm in length (Hamilton & Caselle, 2015).

For the fish surveys, we calculated the number and biomass of sheephead recorded on each transect, and joined these data to the dataset of all unique fish transects. For the invertebrate surveys, we calculated the total number of urchins (summing all three species) and spiny lobsters recorded on each transect, and again joined these data to the dataset of all unique swath transects. Because searches were performed for all species of interest, a value of zero was assumed wherever one of the species was not reported. We estimated California sheephead biomass using length-weight equation for California sheephead b = 0.0144* *l* ^3.04^, where *b* is the biomass in g and *l* is the total length in cm (Hamilton & Caselle, 2015). Next, we summarized these data to average annual abundances per protection category per site (as measured in standard transects), joined the fish and invertebrate data together, and then calculated the average (and standard error) annual abundance for the species of interest across sites from 2002–2023. Thus, each site is equally weighted. Finally, we added a variable called “heatwave”, and assigned its values as “before” (2002–2013), “during” (2014–2016), and “after” (2017–2023), according to the year the data were collected.

There were 81 monitoring sites (32.9%) within fully protected areas, 33 sites (13.4%) within partially protected areas and 132 unprotected sites (53.7 %). Divided by region, there were 120 sites (48.8%) within Southern and 126 sites (51.2%) within Central California. All sites with data we used for analyses are visualized in Supplementary Figure 2.

#### 2.4.2 Regression Models

We hypothesized that higher abundances of sheephead and lobster (mesopredators frequently targeted by fisheries) inside MPAs in Southern California would result in greater predation pressure on sea urchins, thereby decreasing sea urchin kelp herbivory and allowing for greater kelp area and/or faster kelp recovery. We focused on Southern California to examine whether trophic cascades may be a mechanism underlying kelp resilience because only in this region are the main predators of sea urchins directly targeted by fisheries, and therefore benefit from protection in MPAs. To investigate these hypotheses, we used two generalized linear mixed models (GLMMs) to explore the variability in urchin abundances among times and locations. First, we modeled urchin abundances in Central California and Southern California as a function of protection level, period (relative to the MHWs), and interactions between protection and period. Second, we explored whether in Southern California the abundances of California sheephead and spiny lobsters explain variability in urchin abundances (within the conceptual framework of a predator-prey model), allowing for linear and quadratic effects of predator biomass. We modeled these hypotheses separately because the proximal effect of California sheephead predation could mask the effect of protection.

Due to zeros within the sea urchin and sheephead data and the fact that we modeled average urchin count densities, we selected a Tweedie distribution with a log link function for all models. In fitting each model, we estimated the Tweedie power parameter jointly with the model coefficients. Additionally, we fit site-level random intercepts and slopes within both models to account for repeated sampling at each site. The models including random intercepts and slopes were selected based on diagnostic plots of the model residuals, as well as the fact that these models had lower AIC values than those including only random intercepts. Following model fitting, we assessed whether there was evidence of residual temporal autocorrelation in the model by computing the lag-1 autocorrelation on the residuals of each site separately. We found that the average residual autocorrelation among sites was low (0.14). To be sure, we ran both models with and without consideration of a site-level autoregressive order-1 (AR(1)) error structure. No large differences were detected for the models describing the relationship between protection, heatwave period, and urchin abundances; therefore, we chose the simpler model without the autoregressive function. For the predator–prey model, we report both model specifications.

Additionally, we ran three GLMMs to test whether there were greater abundances of spiny lobsters and California sheephead, and greater biomass of California sheephead within protected sites from 2012 onward. We selected a Tweedie distribution with a log link function for all models again and we fit the full models with a site-level AR(1) error structure, and site-level random slopes and intercepts. We then selected model structure on the basis of model fit, removing the random slopes and/or autocorrelation in the error structure on the basis of improvements in model fit. We used the R packages “glmmTMB” to fit our models, “car” to compute Wald Tests of the main effects, and “DHARMa” to assess the model residuals (Brooks et al. 2017; Fox, J. W., 2019; Hartig, F., 2022).

Previous studies have found that MPA age is correlated with increased fish biomass (Claudet et al., 2008; Micheli et al., 2004; Ziegler et al., 2023). To investigate this dynamic, we also used a two-way fixed-effects model to test whether accounting for the year of MPA implementation (spanning from 1973–2012) modified the effect of protection on urchin abundance in Southern California. In this instance, we used an ordinary least-squares estimation with fixed effects for both site and year, and Driscoll-Kraay standard errors. These results are reported in supplementary figure 6.

All data and statistical analyses were carried out in R (version 4.3.1) using R Studio (version 2023.06.0 for windows). All code used for the data preparation, statistics, and figures can be found on the GitHub repository: https://github.com/jkumagai96/Kelp_Forests_and_MPAs

## 3. Results

### 3.1 Resilience of kelp within MPAs to MHWs

Landsat data reveal that during the 2014–2016 MHW there was an average loss of 46.4% of kelp canopy area relative to the historic baseline in Southern California. In 2017–2021, there was some recovery, but coverage remained 39.1% below the baseline. In Central California kelp cover loss during the MHWs with respect to the baseline was lower than in Southern California, i.e. 13.3%, but there was no recovery during the 2017-2021 period, kelp forest loss increased to 24.8% with respect to the historic baseline.(Figure 2).

**Figure 2:**
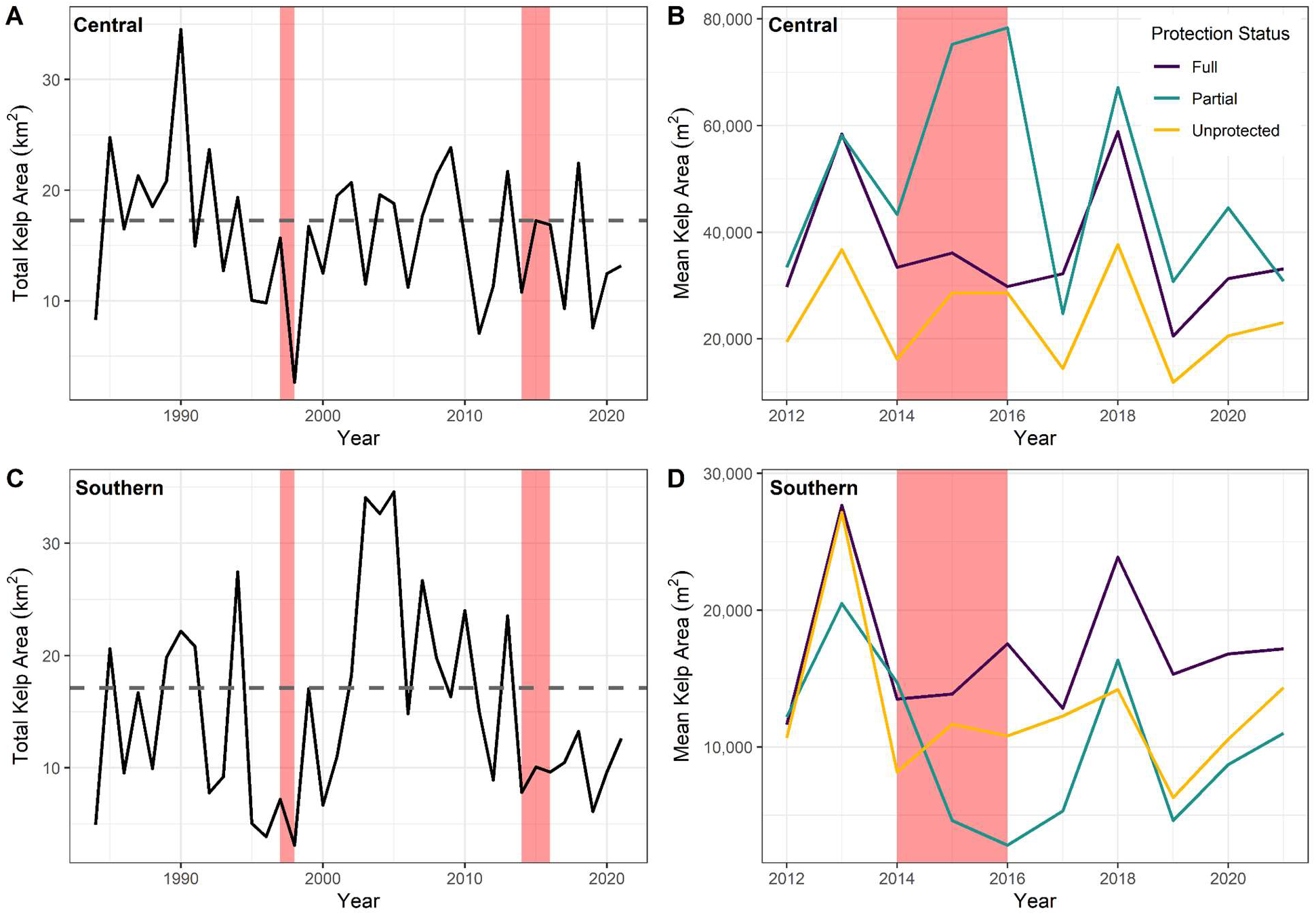
Kelp area through time for the study area. The left column reports total kelp area (km^2^) within Central (A), and Southern California (C), with the mean baseline kelp area between 1984–2013 represented as a horizontal dashed line. The right column reports mean kelp area in m^2^ per 1-km^2^ pixel by protection category from 2012–2021, to include all MPAs established in southern and central California, with kelp from fully protected areas in purple, partially protected areas in turquoise, and unprotected areas in yellow for Central (B), and Southern California (D). Note that the axes are not held constant and MHWs (the 1997-98 extreme ENSO event and 2014-16 MHW) are denoted using transparent red rectangles.

Both during and after the MHWs, there was significantly higher relative kelp area within fully protected areas than unprotected areas (Figure 3, *p* < 0.005), while there were no significant differences in kelp area between partially protected and unprotected areas (Figure 3 A-B; Supplementary Table 2). However, this overall pattern is driven by responses in southern California. When analyzed by region, the only significant differences in relative kelp area in Central California were between partially protected and unprotected areas during the MHW (Figure 3; Supplementary Table 2), with more kelp within partially protected areas during the MHW (Figure 2B & Figure 3C). In Southern California, there was significantly higher resistance to, and recovery from, MHWs within fully protected areas compared to partially protected and unprotected areas (*p* < .05, Figure 2E-F). Importantly, we found no significant difference between relative kelp area within partially protected areas and unprotected areas in Southern California. Based on this evidence, fully protected areas appear to confer resilience to MHWs, both in terms of resistance and recovery, depending on the region.

**Figure 3:**
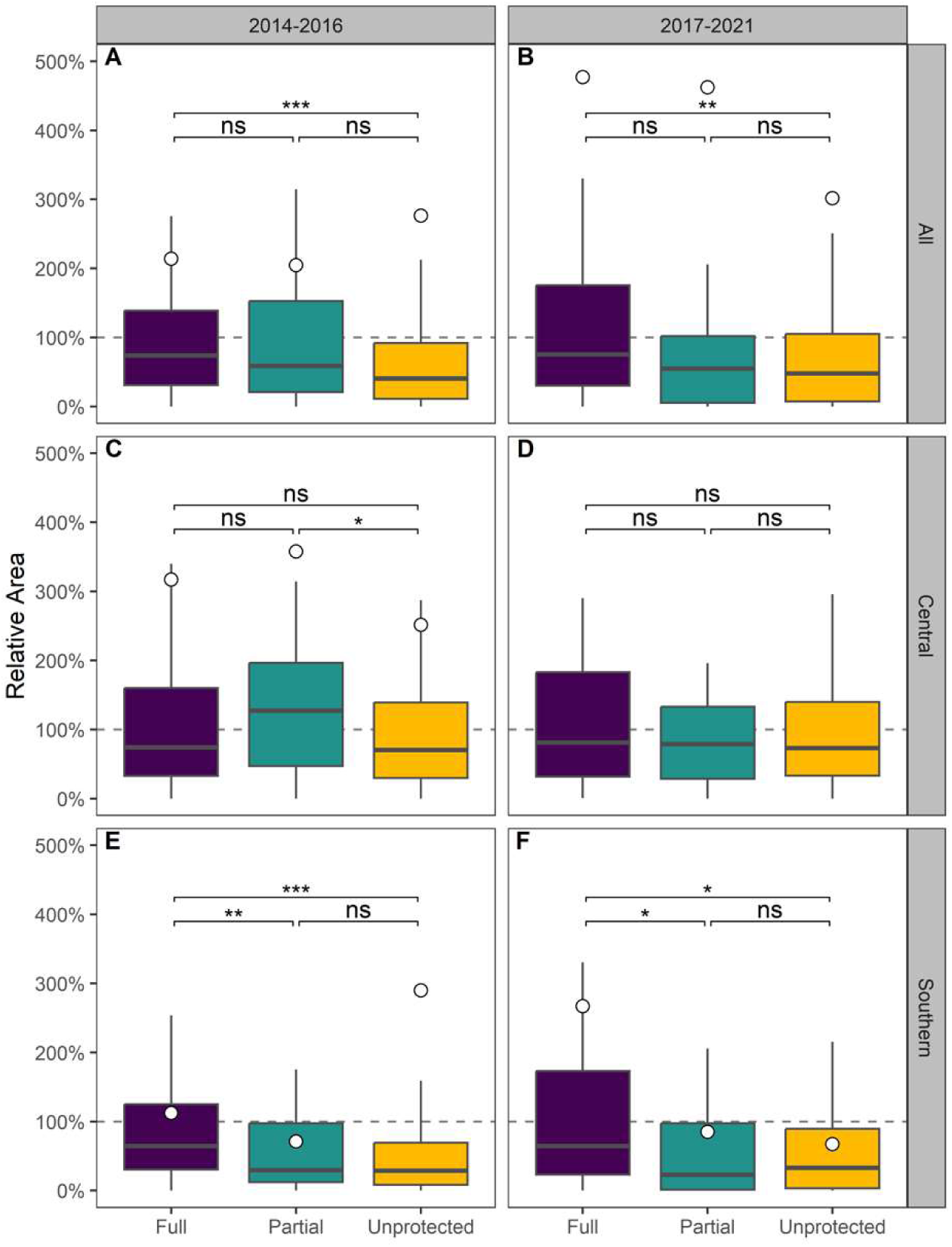
Resilience of kelp forests during (2014–2016) and after the MHWs (2017–2021) Boxplots of relative area of kelp (averaged annual kelp area relative to the historic baseline area within each pixel) are reported within fully protected, partially protected, and unprotected areas for (A-B) all regions, (C-D) Central California, and (E-F) Southern California. White points represent averages. Average points in Central California in 2017–2021 are outside the plot extent and not visualized (see Supplementary Table 5) and outliers are also removed from the plot for ease of visualization. Pseudo p-values were computed via Bonferroni-corrected permutation analyses; non-significant group differences are indicated with “ns” while significant comparisons (after Bonferroni correction) are denoted with asterisks — *p* < 0.05 (*), < 0.01 (**), and < 0.001 (***).

When assessing the impact of MPA age on these results in Southern California, we found that kelp forests within fully protected areas consistently had significantly higher resistance to the MHWs independent of MPA age, although the effect was stronger in MPAs established before 2007 compared to the younger MPAs (Supplementary Figure 5). However, recovery was indistinguishable between new and old MPAs, albeit that new MPAs exhibited significantly higher relative area of kelp in fully protected areas compared to partially protected areas (Supplementary Figure 5). Additionally, when testing how sensitive our results were to high values in percent recovery, we found that kelp forests within fully protected areas consistently had higher resistance to the MHWs than unprotected areas, although recovery was sensitive to these high values (Supplementary Table 6). Taken together, these results could be biased if MPAs had been non-randomly placed in habitat more favorable to kelp recovery. A principal component analysis revealed no evidence for difference in environmental variables (i.e. temperature, depth, marine heatwave intensity) between protection categories from before (2013), during (2015), and after (2019) the 2014–2016 MHWs, suggesting that protection status is not correlated with pre-existing resilience potential (Supplementary Figure 1).

### 3.3 Mechanism of resilience: trophic cascades

In Central California, urchin abundances significantly increased overall from 2014**–**2023 in all protection categories (Figure 4, ꭓ^2^ = 684, df = 2, *p* <0.0001). Average urchin abundance across all protection categories was only 2.61 ± 0.58 per transect before the MHWs but increased to 123 ± 28 per transect during the MHWs and 371 ± 89 per transect after the MHWs. There was no significant interaction between protection category and heatwave period, suggesting that protection status had no effect on urchin abundance and their population increase during and after the MHWs.

**Figure 4:**
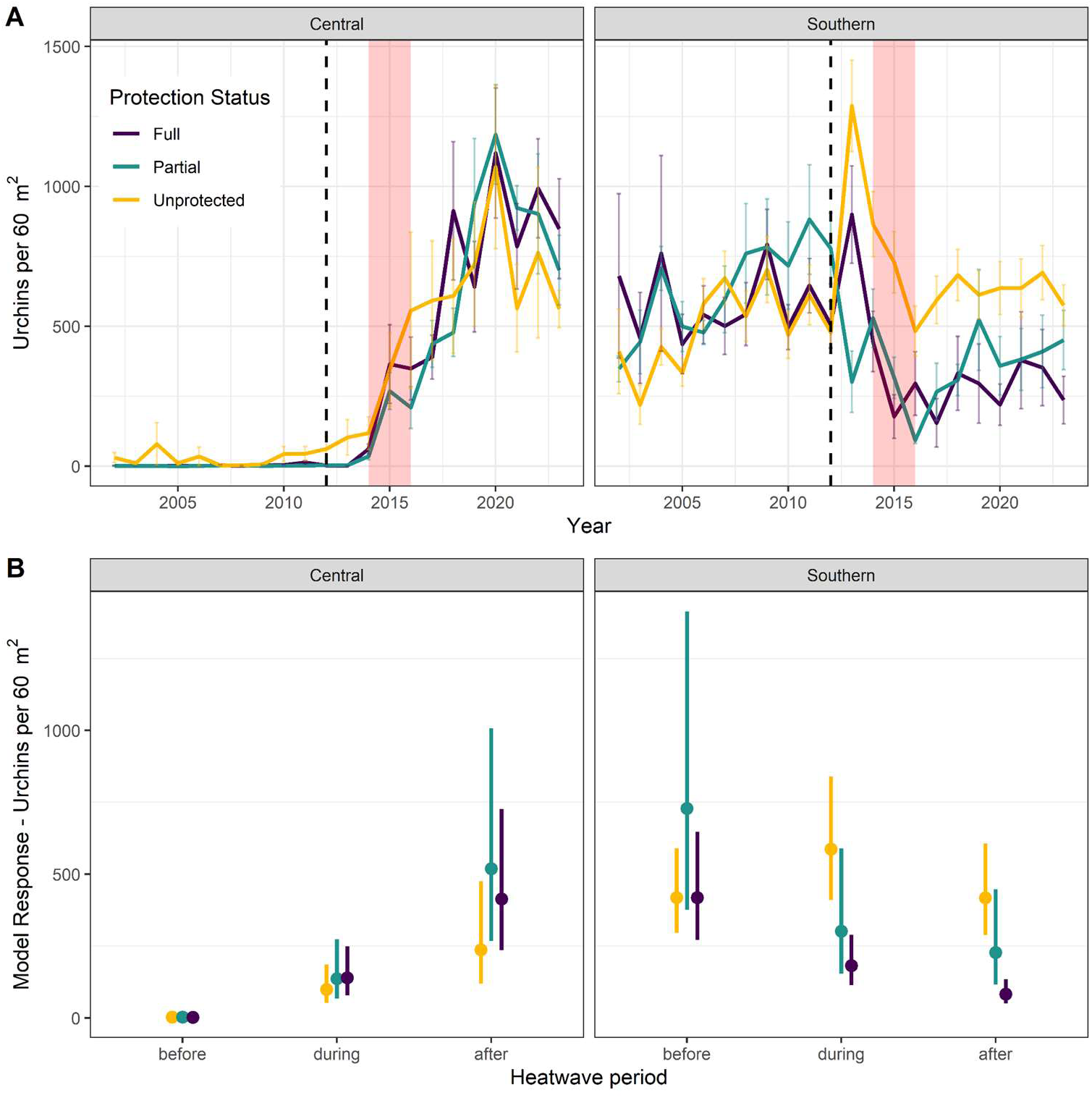
Urchin dynamics through time by level of protection in Central and Southern California. (A) Mean urchin abundances per site and level of protection (number of individuals per standard 60 m^2^ transect, *N* = 121 and 94 sites for Central and Southern California respectively) from subtidal data for the period 2002–2021 from PISCO. All urchin species were combined. The dashed line at 2012 represents the implementation of the last MPAs under the Marine Life Protection Act. Error bars represent standard errors. Data before 2012 include sites that were protected at that time or would become protected in 2012. The 2014–2016 MHWs are depicted in transparent red. (B) Variation in urchin densities across protection levels (full, partial, and unprotected) and heatwave periods (before, during, and after) for both regions. Points and line ranges represent estimates and confidence intervals (respectively) for mean urchin density from a Tweedie GLMM. The vertical lines in panel B represent 95% confidence intervals.

In Southern California, overall urchin density in unprotected areas (418 ± 73 per transect (60m^2^) mean ± SE)) before the MHWs was not significantly different from that in partially (729 ± 246, *p* = 0.30) or fully protected (419 ± 93 per transect, *p* = 1) areas before the MHWs. However, we found that the difference in urchin abundance between protection categories varied through time (ꭓ^2^ = 84, df = 4, p < 0.0001). In contrast with Central California, urchin abundance was significantly lower in fully protected areas than in unprotected areas both during (*p* < 0.0001) and after (*p* < 0.0001) the MHWs. Urchin abundances also declined in partially protected areas during (302 ± 103 per transect) and after (228 ± 78 per transect, *p* = 0.039) the MHWs, but these abundances were only statistically significantly different from those for unprotected areas after the heatwave (Figure 4). There were significantly fewer urchins in fully protected areas (83 ± 21 per transect) compared to partially protected areas (228 ± 79 per transect) after the MHWs (*p* = 0.039). Using a two-way fixed-effects model, we found that urchin abundances declined with MPA age, particularly in fully protected areas (Figure S6). Taken together, these results indicate that the difference in urchin abundances between unprotected and protected areas increased during and after the MHWs.

After the full implementation of the Marine Life Protection Act and establishment of all MPAs, completed in 2012, there was a significant increase in spiny lobster abundance within fully protected sites compared to both unprotected (*p <* 0.0001) and partially protected sites (Figure 5A, *p* = 0.0055) in Southern California. Sheephead abundance increased at all sites during the MHW (Figure 5B). Surprisingly, after the MHW, there were significantly higher abundances of California sheephead within partially protected sites compared to fully protected and unprotected sites (Figure 5B, *p* = 0.0052). When assessing differences in California sheephead biomass, there was significantly higher biomass within both fully (*p* = 0.011) and partially protected sites (*p* = 0.002) compared to unprotected sites (Figure 5C). The different patterns observed between abundance and biomass trends within fully protected sites suggest larger sheephead sizes in fully protected sites.

**Figure 5:**
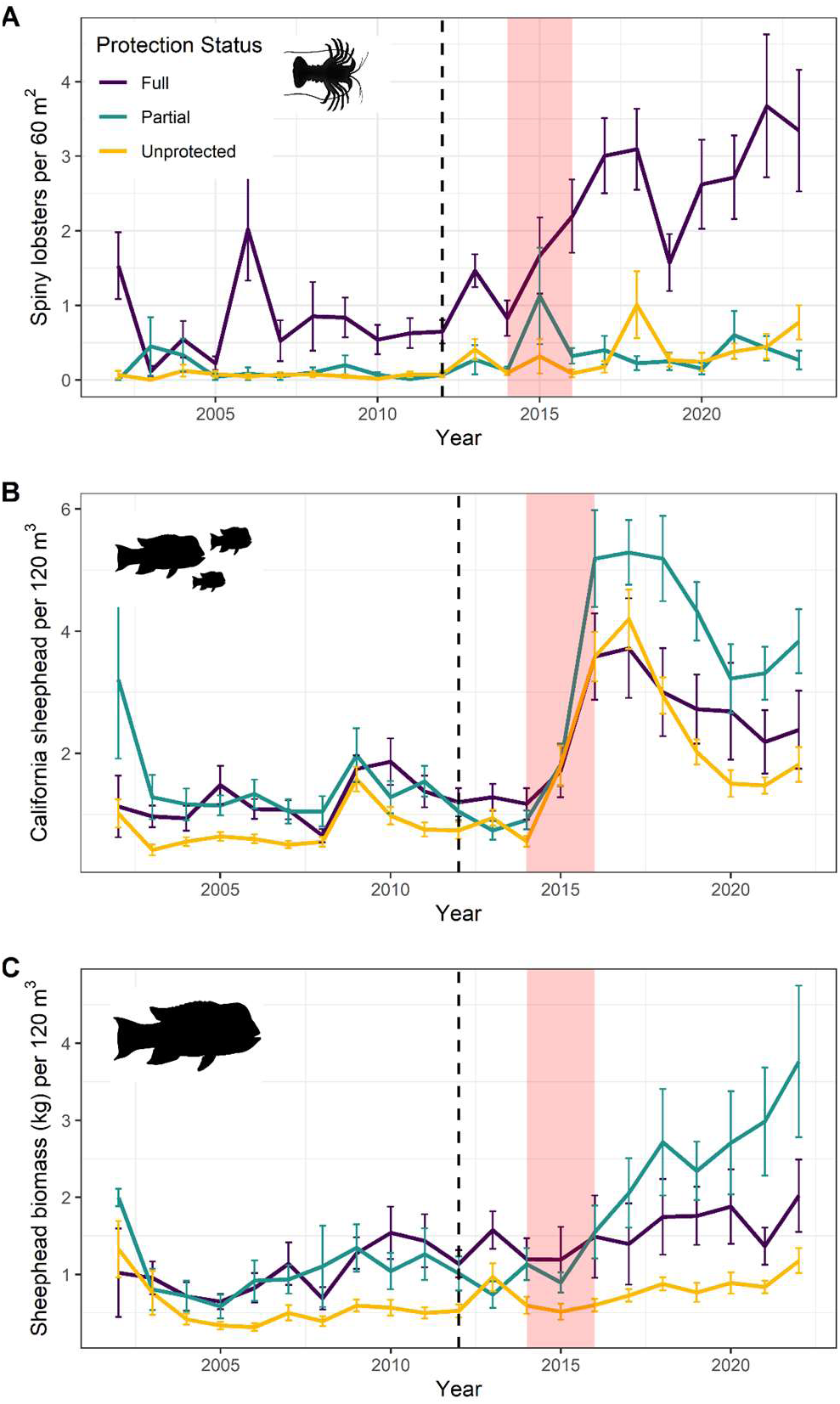
Average abundances of urchin predator population size per site and level of protection for Southern California from 2002–2023. (A) Mean abundances of spiny lobsters per site (number of individuals/60m^2^), (B) Mean abundance of California sheephead per site (number of individuals/120m^3^), (C) Mean biomass of California sheephead (g/120m^3^). The dashed line at 2012 represents the implementation of the last marine protected areas under the Marine Life Protection Act. Error bars represent standard errors. Data before 2012 includes sites that were protected or would become protected in 2012. Data within fully protected areas are in purple, partially protected areas in turquoise, and unprotected areas in yellow. The MHWs in 2014–2016 are depicted in transparent red.

When assessing predators’ relationship to urchins, we found that the abundances of California sheephead and spiny lobster were negatively correlated with abundances of urchins (Figure 6). When accounting for temporal autocorrelation, abundances of California sheephead were negatively correlated with urchins, but spiny lobsters were not (Figure 6, residuals in Supplementary figures 7-10). Regardless of model choice, the relationship between the abundances of California sheephead and urchins was consistently negative and significant.

**Figure 6:**
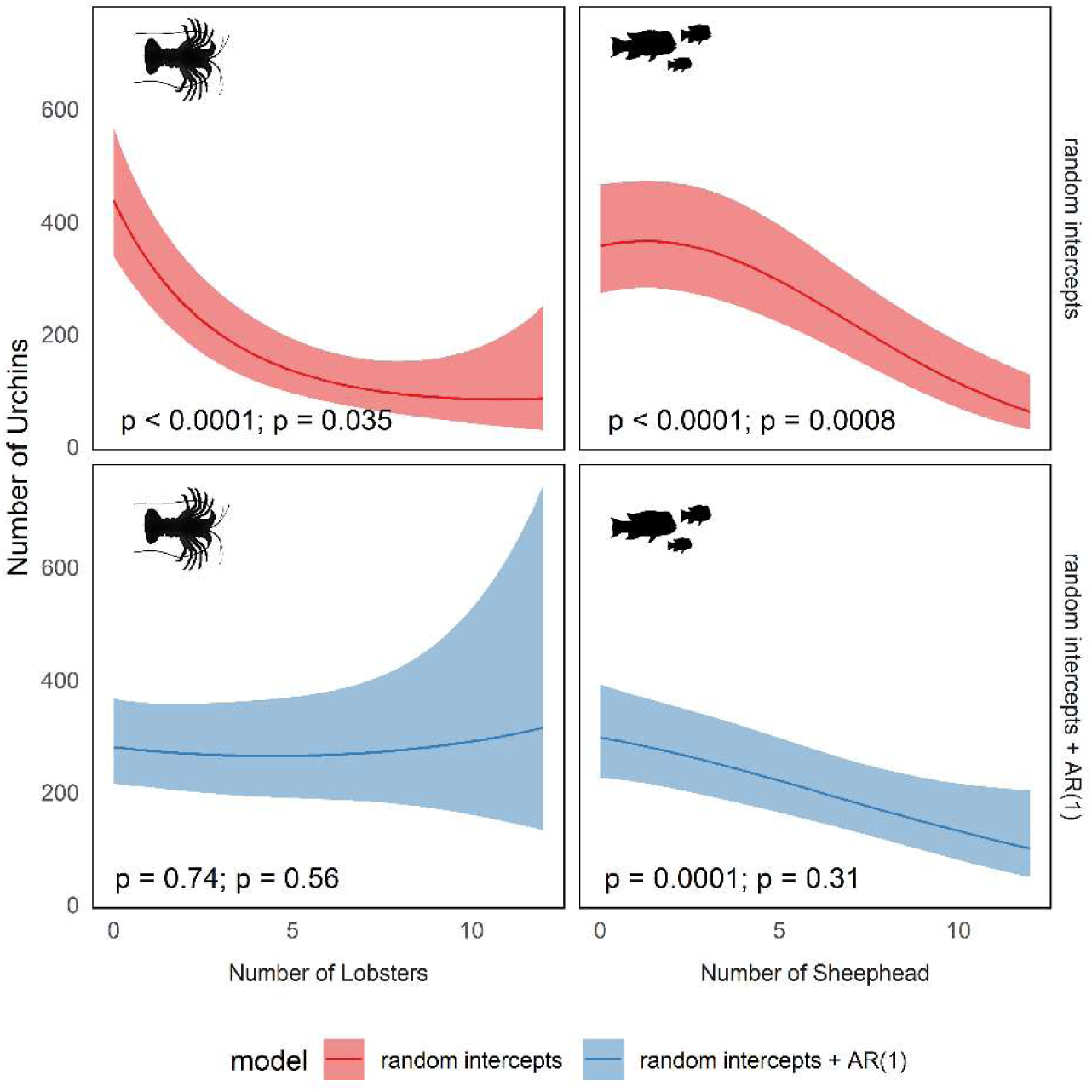
Partial effects plots from two models of average urchin abundances per site in Southern California as a function of California sheephead and spiny lobster abundances. One model includes an autoregressive correlation structure (blue), and the other does not (red). On each panel there are two significance values, the left-most *p* value corresponds to the log-linear effects, while the right-most *p* value corresponds to the log-quadratic effects. Plots for model residuals can be found in the supplementary (Supplementary Figure 6 and 7).

## 4. Discussion

This study provides empirical evidence that fully protected MPAs can promote the resilience of kelp forests to climate impacts when protection has a positive effect on natural predators of sea-urchins. Full protection improved both kelp resistance to, and recovery from, extreme MHWs. This pattern emerged for Southern California, suggesting that the current network of MPAs offers resilience to climate change impacts, but only when and where MPAs successfully protect urchin predators. In Central California, where the main urchin predators have been depleted by a disease outbreak (sunflower sea stars) or are protected statewide and therefore not directly influenced by MPA status (sea otters), sea urchins increased dramatically during and after the MHW, across both protected and unprotected sites. In contrast, in Southern California, protected areas had significantly greater abundances of urchin predators and fewer urchins within both partially and fully protected MPAs during and after the 2014–2016 MHWs. These results lend support to the role of trophic cascades as a mechanism for ecological resilience, and fully protected MPAs as a climate adaptation tool.

Our findings provide evidence that trophic cascades are a mechanistic path through which MPAs provide climate resilience to kelp forest ecosystems; however, these benefits are highly context-dependent and vary regionally. Multiple studies have shown that fully protected MPAs increase the biomass and abundance of the predators of urchins (Caselle et al., 2015; Hamilton & Caselle, 2015; Lenihan et al., 2022), which exerts a top-down control on urchin populations, thereby supporting stability and resilience of kelp populations (Ling et al., 2009; Peleg et al., 2023). Here, we show that this mechanism also applies under climate impacts because we observed that there were fewer sea urchins, less loss of kelp and greater recovery of kelp populations inside fully protected MPAs during and after the 2014–2016 MHW in Southern California. Corroborating this interpretation of our results, we found that urchin abundances were negatively correlated with those of spiny lobster and California sheephead. These results suggest that the recovery of urchin predators within protected areas from overfishing is likely controlling urchin populations and potentially behavior, thus preventing overgrazing and allowing kelp to recover faster from disturbances than in unprotected areas.

In Central California, we found no measurable effect of protection status on kelp resistance and recovery, likely because spatial protection does not confer additional benefits to the main mesopredators of urchins in the region—sea otters and sunflower sea stars—whose dynamics are largely independent of fishing effort and, as a consequence, protection status. Sea otters are federally protected and have not been actively hunted for over a century, thus benefiting from protection throughout their range. Further, sea urchin abundance started to increase exponentially both inside and outside protected areas following the mass mortality of sea stars due to the outbreak of sea star wasting disease in 2013–2015, which led this sea urchin predator to near extinction (Harvell et al., 2019; Montecino-Latorre et al., 2016; Rogers-Bennett & Catton, 2019). We assume that the level of protection has no influence on recovery of sea stars, as this species is not being actively harvested and has yet to recover. These observations illuminate how non-spatial policies, such as species-specific interventions (i.e. federal protection conferred over sea otters, and possibly the proposed active restoration of depleted seastar populations through captive breeding and outplants) may also promote some degree of ecosystem resilience.

Our results are consistent with and expand on other studies in the region, emphasizing evidence for trophic cascades—preserved by MPAs—as the mechanism separating healthy kelp forests from urchin barrens. For example, trophic cascades were found to enhance macroalgae abundances in MPAs in the northern Channel Islands a year after the MHWs (Eisaguirre et al., 2020). However, another study in the Channel Islands found contrasting evidence: there was an increase of urchins within protected areas, in part due to the release of red urchins from fishing pressure within MPAs, which outweighed any effect of trophic cascades (Malakhoff & Miller, 2021), though the authors of this study did not consider the response of urchins to the MHWs. In comparison, we found fewer urchins within MPAs, but only during and after the MHWs. Notably, when we took into consideration the year of establishment for the MPAs, we found that protection led to fewer urchins in Southern California through time (Supplementary figure 6). Therefore, by expanding the spatial and temporal scale of analysis, our results reconcile previously contrasting conclusions.

Our work is also subject to some limitations. First, the long-term dataset of kelp area tracks only the area of kelp at the surface; we have no remote sensing information on subsurface giant kelp. Additionally, kelp area is an estimate from satellite imagery which may add some sources of error (Alix-Garcia & Millimet, 2022). However, ongoing methodological improvements have addressed most detection gaps (see Bell et al., 2020 for more detail). For the subtidal data, while we have size structure information for California sheephead that allowed us to evaluate biomass, such data are not available for spiny lobsters as it is difficult to measure their size in the field. Larger individual biomass of overexploited species is expected inside fully protected MPAs, and these larger animals are usually more efficient at consuming larger urchins (Hamilton & Caselle, 2015). Having such estimates for spiny lobsters in this study could help us to further understand the role of spiny lobsters in trophic cascades, although spillover effects of lobsters in both abundance and biomass have been demonstrated previously (Lenihan et al., 2022). Moreover, we did not include in our analyses other smaller species, such as crabs, which may benefit from MPAs and influence urchin populations by feeding on their juveniles (Clemente et al., 2013). We excluded these species because of the current limited understanding of their role as urchin predators. Finally, we were not able to explore evidence for trophic cascades within Central California as there is no population data for otters at the same scale and resolution of the PISCO data.

Besides trophic interactions, there are additional potential reasons why spatial protection in Central California was not associated with increased climate resilience for kelp forests. First, this region was less impacted by the MHWs and had an overall better recovery from the MHWs than Southern California. Notably, on average (not considering protection status) kelp area remained on average 1.5 - 3 times higher in central than southern California during and after the MHW (Figure 2). It is no surprise that level of protection had no effect in Central California because, regardless of protection, giant kelp forests were not as impacted from the MHWs, although they had a steady decline after the heatwaves. Giant kelp in Central California were more resilient during the MHWs likely because temperature extremes during the MHWs were often below the thermal tolerance limit of giant kelp. In addition, large areas of Central California are less accessible to people and therefore less impacted by human activities, including fishing (Free et al., 2023), than in Southern California, and because density of the remaining urchin predators (federally protected sea otters) is largely uncorrelated with protection status. Our results are in general agreement with previous studies that also found limited contribution of MPAs to climate resilience for kelp forests communities in Central California (J. G. Smith et al., 2023). These findings suggest that it is a priority to assess the benefits of MPAs for providing climate resilience in regions that are more impacted by climate change and human activities. For example, research is needed to evaluate whether MPAs or other management strategies provide similar benefits near the distribution limit of giant kelp in Baja California Sur, Mexico. Our study casts new light on differences in climate resilience between two regions in California and, most importantly, highlights the importance of the local ecological context in determining whether MPAs can be expected to buffer climate extremes.

Our findings have important implications for evaluating the benefits that MPAs can confer in terms of mitigating the impacts of climate change, and also for informing approaches to climate-smart management and establishment of new MPAs (Arafeh-Dalmau et al., 2023) as nations make progress toward protecting 30% of the oceans by 2030 while adapting to climate change (Convention of Biological Diversity, 2022). Understanding which mechanisms provide climate resilience at different levels of biological organization (species, population, and ecosystem), and at local to regional scale, is crucial to inform realistic expectations of the benefits of resilience to climate change that MPAs or other management options may provide. There is a need for deeper understanding of the local biogeography, environmental conditions, and management strategies that drive ecosystem resilience to understand where placing MPAs may increase climate resilience. Furthermore, such understanding will require investment in long term monitoring and standardized metrics to define and measure ecological resilience to evaluate the conditions under which MPAs confer resilience to climate impacts.

The most important implication of our findings is that protection of top predators confers benefits that propagate through the ecosystem, boosting resilience to and recovery from acute impacts of climate change. While this goal often underpins the establishment of MPAs, its effectiveness in providing climate resilience is seldom supported by empirical evidence. Of course, additional research is required to assess the generality of our findings, but they provide a strong initial motivation to carefully manage fishing pressure in the coastal zone as climate extremes become more frequent and intense (Oliver et al., 2018; Schoeman et al., 2023). Protected areas offer many benefits from preventing continued destruction of habitats (including blue carbon ecosystems such as seagrass and mangroves), increasing food security, and increasing resilience to climate shocks and environmental variability, ultimately increasing overall ecosystem resilience (Aburto-Oropeza et al., 2011; Jacquemont et al., 2022; Miteva et al., 2015; Selig & Bruno, 2010). However, protected areas are not a panacea to the ongoing and projected impacts of climate change. In particular, our results of context-dependent roles of MPAs in providing climate resilience highlights the urgency to carefully consider what and where additional measures are needed, such as the protection of wide ranging top predators to the active restoration of habitat and critical species interactions. Crucially, the root causes of climate change and global biodiversity loss must be urgently addressed before the efficacy of our adaptation tools is lost (Mills et al., 2023).

## Acknowledgements

We want to thank the Santa Barbara Coastal LTER, PISCO, NOAA, and the many other data providers for sharing their data openly so our mutual understanding of these systems can continue to advance. In addition, thank you, Joel Erberich, for the many discussions that helped to form the paper in its current form. Thank you, Marty Freeland, for the black and white organism figures which help to clearly communicate our science. Also, thank you Ryan O’Connor for the helpful feedback on data visualizations. TB and the kelp canopy and environmental dataset was funded by the NASA Biodiversity and Ecological Conservation award #80NSSC22K0169. NA-D, FM, and KC acknowledge funding from the Lenfest Ocean Program (ID Number: 00036969) to explore climate resilience in marine protected areas. Finally, we acknowledge funding from the NSF Award #2108566 - DISES: Pathways and constraints to adaptation in coastal social-environmental systems.

## Data Availability

The data that support the findings of this study will be openly available in Zenodo at —— [will be updated if accepted.

These data were derived from the following resources available in the public domain: the Kelpwatch SBC LTER dataset (https://doi.org/10.6073/pasta/c40db2c8629cfa3fbe80fdc9e086a9aa), the marine protected area dataset (https://marineprotectedareas.noaa.gov/dataanalysis/mpainventory/), and an updated version of the currently publicly available PISCO data (https://data.piscoweb.org/metacatui/view/doi:10.6085/AA/PISCO_kelpforest.1.6).

## Author Contributions

**Joy A. Kumagai** – Conceptualization, data processing and analysis, investigation, methodology, visualizing, writing – original draft. **Maurice Goodman** – Data processing and analysis, methodology, visualizing, writing – review and editing. **Juan Carlos Villaseñor-Derbez** – Data processing and analysis, methodology, writing – review and editing. **David S Schoeman** – conceptualization, investigation, writing – review and editing. **Kyle Cavanuagh** – conceptualization, writing – review and editing. **Tom Bell** – conceptualization, writing – review and editing. **Fiorenza Micheli** – conceptualization, investigation, writing – review and editing. **Giulio De Leo** – investigation, methodology, writing – review and editing. **Nur Arafeh-Dalmau** – conceptualization, data processing and analysis, investigation, methodology, writing – review and editing.

