## Supplementary Information for "Marine protected areas promote resilience of kelp forests to marine heatwaves by preserving trophic cascades"

#### 1) Principal Component Analysis to evaluate similarities in environmental conditions

*Methods*

Protected areas may be preferentially placed in the most pristine, productive, or biodiverse habitats. To test this assumption, we evaluated whether MPAs in our study area are in more productive waters, to assess whether they exhibited higher resistance and recovery simply because they are in more productive areas, and not because of their protection level. We tested whether this was the case by gathering biophysical and socioeconomic parameters (described below), which were used to perform a principal component analysis to explore clustering of points in relation to protection. Specifically, we obtained yearly estimates of surface nitrate concentrations, temperature, maximum significant wave height, and depth which are known variables to influence the dynamics of kelp forests. This information was obtained from the Santa Barbara Coastal Long Term Ecological Research project (Bell, T.W., 2023).

Then we identified MHW and cold spells using the "heatwaveR" package (Schlegel and Smit, 2018). Daily sea surface temperature was downloaded from NOAA's 0.25-degree OISST dataset from 1983 to 2021 (https://doi.org/10.25921/RE9P-PT57). MHW events and cold spells are defined as periods where sea surface temperatures are above the 90$^{th}$ percentile and below the 10$^{th}$ percentile, respectively, based on a 30-year baseline climatology and last for at least 5 consecutive days (Hobday et al., 2016). For this analysis we used climatology from 1983 to 2012. We then estimated the yearly cumulative MHW and cold spells intensities (°C days). Cumulative intensities are good indicators of the exposure of species to warm or cold anomalies in a given year (Arafeh-Dalmau et al., 2023; Oliver et al., 2019).

To estimate the potential influence of human presence along the coastline, we calculated a metric called the human gravity index (Cinner et al., 2018). The human gravity index (HGI) is a useful proxy to understand the level of exposure to human activity a given kelp patch experiences, as it combines the distance to and population size of nearby human settlements. We use publicly available data on the location and size of human settlements for Mexico (2020 census conducted by INEGI, available here: https://www.inegi.org.mx/programas/ccpv/2020/#datos_abiertos) and the United States (2020 American Community Survey from the United States Census Bureau, available here: https://www.inegi.org.mx/app/descarga/ficha.html?tit=326108&ag=0&f=csv) and retain only locations that occur within 50km of the coast (Supplementary Figure 12). We then calculate the HGI for each of the 3,162 pixels with kelp presence with the following equation.

$$HGI_i = \sum_{j=0}^{J} \frac{P_j}{{D_{ij}}^2} \quad (1)$$

Where, $HGI_i$ is the human gravity index of kelp pixel i summed across all human settlements J that fall within 50 km of the kelp pixel, $P_j$ denotes the population size of human settlement j, and $D_{ij}$ is the distance between kelp pixel i and human settlement j.

Finally, we rasterized all the spatial environmental variables to the same 1-km grids of the kelp data, exported the rasters as CSVs, and joined the resulting datasets with human gravity and region data. The MHW and cold spell data had a lower resolution and did not have values close to the coastline for 11.5% of the kelp data. For these kelp area values we used the value of the closest pixel of the MHW and cold spell data.

We used the R package "factoextra" (Kassambara and Mundt, 2020) to run a principal component analysis on the continuous environmental variables (See Supplementary Table 1), except for kelp area. We used the log of human gravity and assigned the smallest non-zero value to any zeros of human gravity. Within the function, "prcomp", we centered and scaled all variables.

*Results and Interpretation*

If MPAs were placed in habitat more favorable to kelp recovery to begin with, differences in resistance to, or recovery from, MHWs allegedly attributed to protection levels might instead be a consequence of different environmental drivers rather than protection status. We examined variation in environmental variables through time and across MPAs and unprotected areas and found that there was no difference in the continuous environmental variables between protection categories from before (2013), during (2015), and after (2019) the 2014–2016 MHWs (Supplemental Figure 1A, 1D, 1G). Therefore, our findings are unlikely to be confounded by MPAs being implemented in healthier or more productive environments (or vice versa). As a positive control, we also visualized the first two principal component axes by region. The principal component analysis confirmed that Southern and Central California are indeed characterized by different environmental conditions in each year (Supplemental Figure 1B, 1E, 1H).

#### 2) Figures

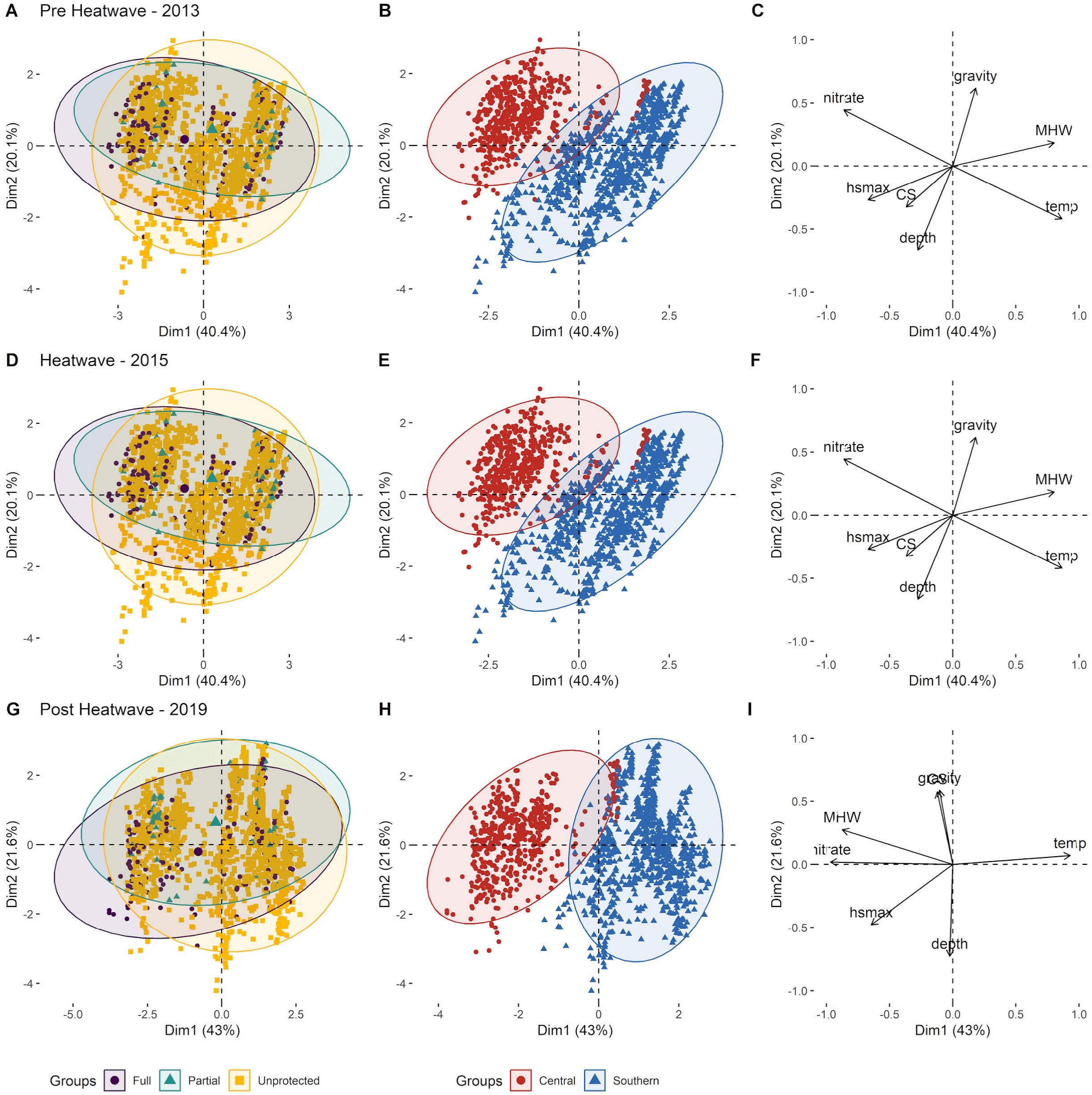


**Supplementary Figure 1: Principal component analyses for all continuous environmental variables considered within the study for before (2013), during (2015), and after (2019) the MHWs.** A, D, and G visualize the principal components by protection level. Data for fully protected areas are purple circles, partially protected areas are turquoise squares, and unprotected areas are yellow triangles. B, E, and H, visualize the principal components by region. Red circles represent Central California, while blue triangles represent Southern California. C, F, and I represent the vectors mapped onto the same axes, with direction and length symbolizing direction and strengths of relationships between variables (note that the scale of panels C, F, and I differs from the other panels). The ellipsoids represent 95% quantiles of the data assuming a bivariate normal distribution.

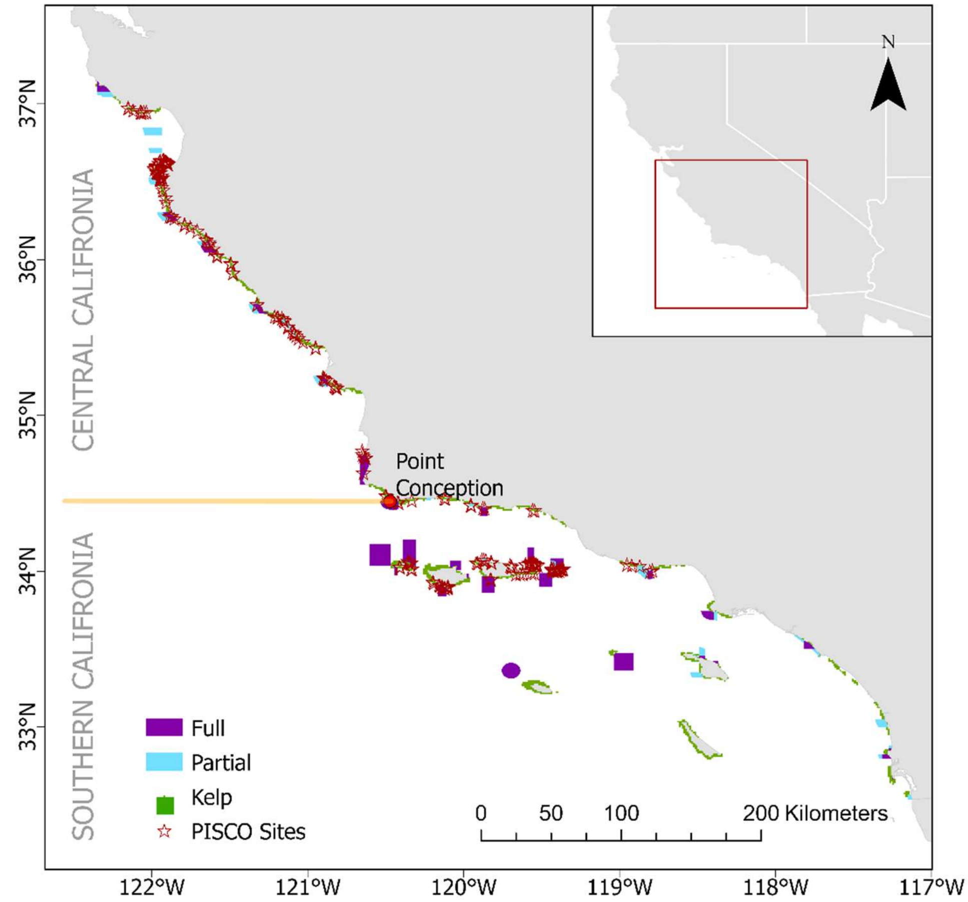


**Supplementary Figure 2:** Study area with locations of PISCO sites included in our study.

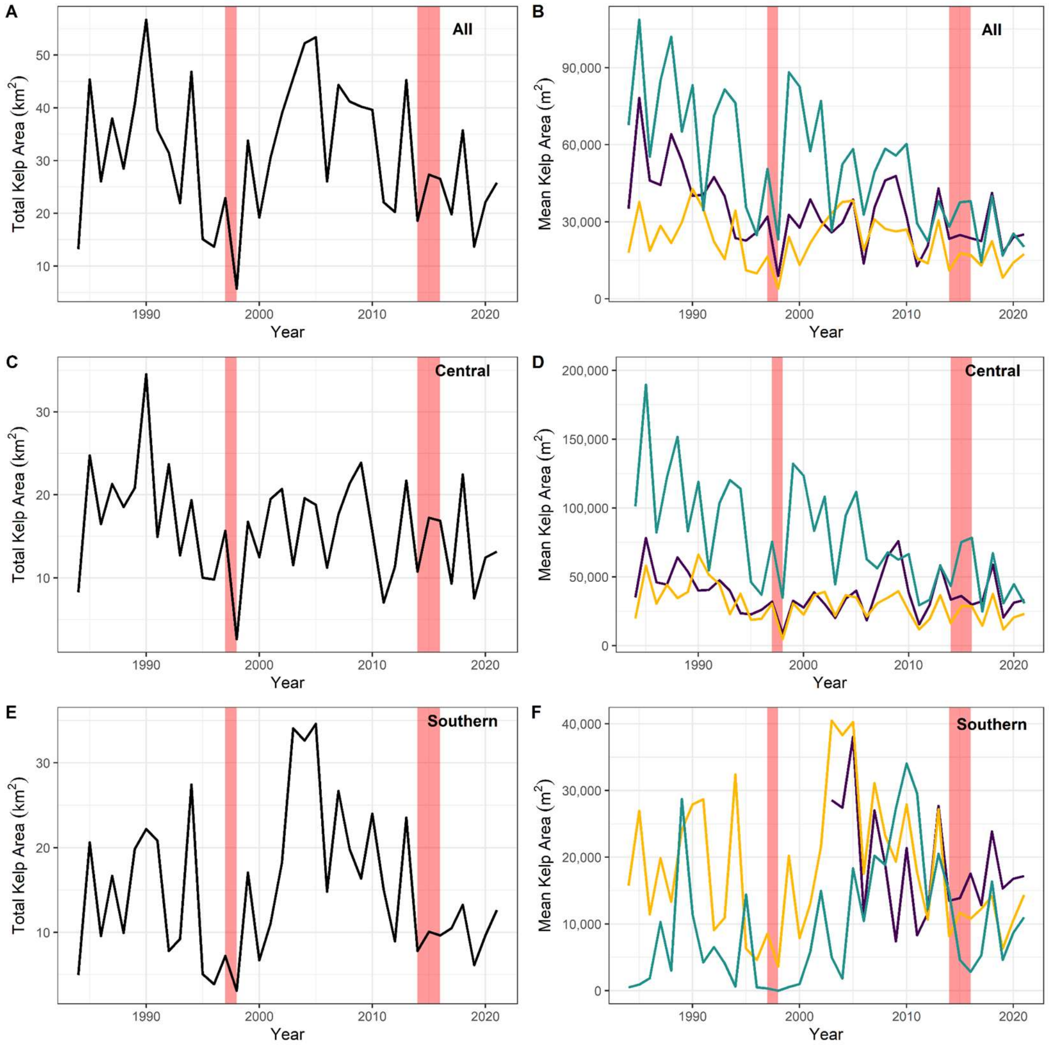


**Supplementary Figure 3:** Kelp area through time for the study area with heat waves denoted in transparent red. The left column reports total kelp area [$km^2$] within Central and Southern California (a), Central (c), and Southern California (e). The right column reports mean kelp area in $m^2$ per 1-km pixel by protection category, with kelp from fully protected areas in purple, partially protected areas in turquoise, and unprotected areas in yellow for both regions together (b), Central (d), and Southern California (f). Note that the axes are not held constant. Before an area is established as protected, the kelp within that area would be considered "unprotected".

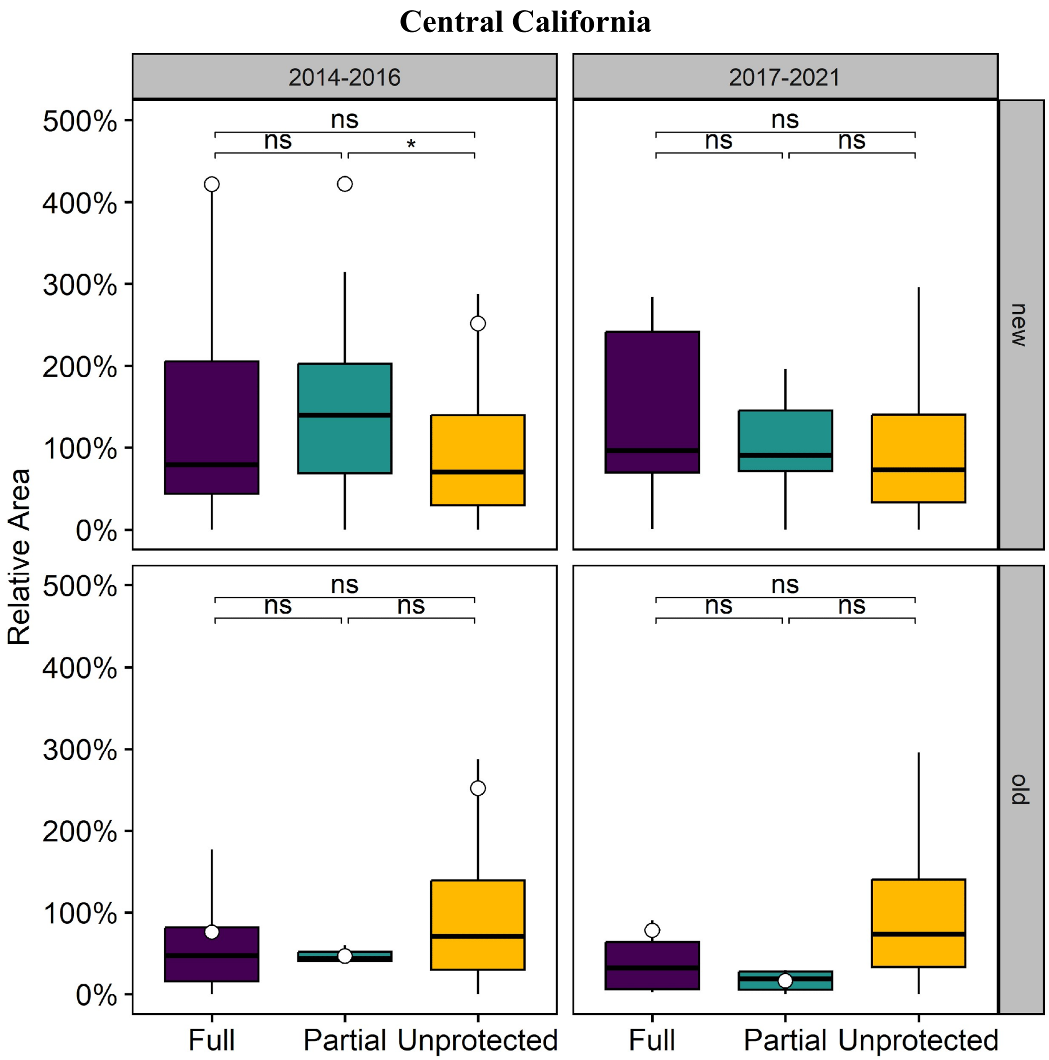


**Supplementary Figure 4**: Mean kelp canopy area relative to the historic baseline for each pixel during (2014–2016) and after the MHWs (2017–2021) in Central California within fully protected, partially protected, and unprotected areas considering only MPAs established after 2007 (top row), and MPAs established before and during 2007 (bottom row). White points represent averages per group. Average points in “new MPAs” in 2017–2021 are outside the plot extent and not visualized. Outliers are also removed from the plot for ease of visualization. Pseudo p-values were computed via Bonferroni-corrected permutation analyses; non-significant group differences are indicated with “ns” while significant comparisons are denoted with asterisks – $p < 0.05$ (*), < 0.01 (**), and < 0.001 (***).

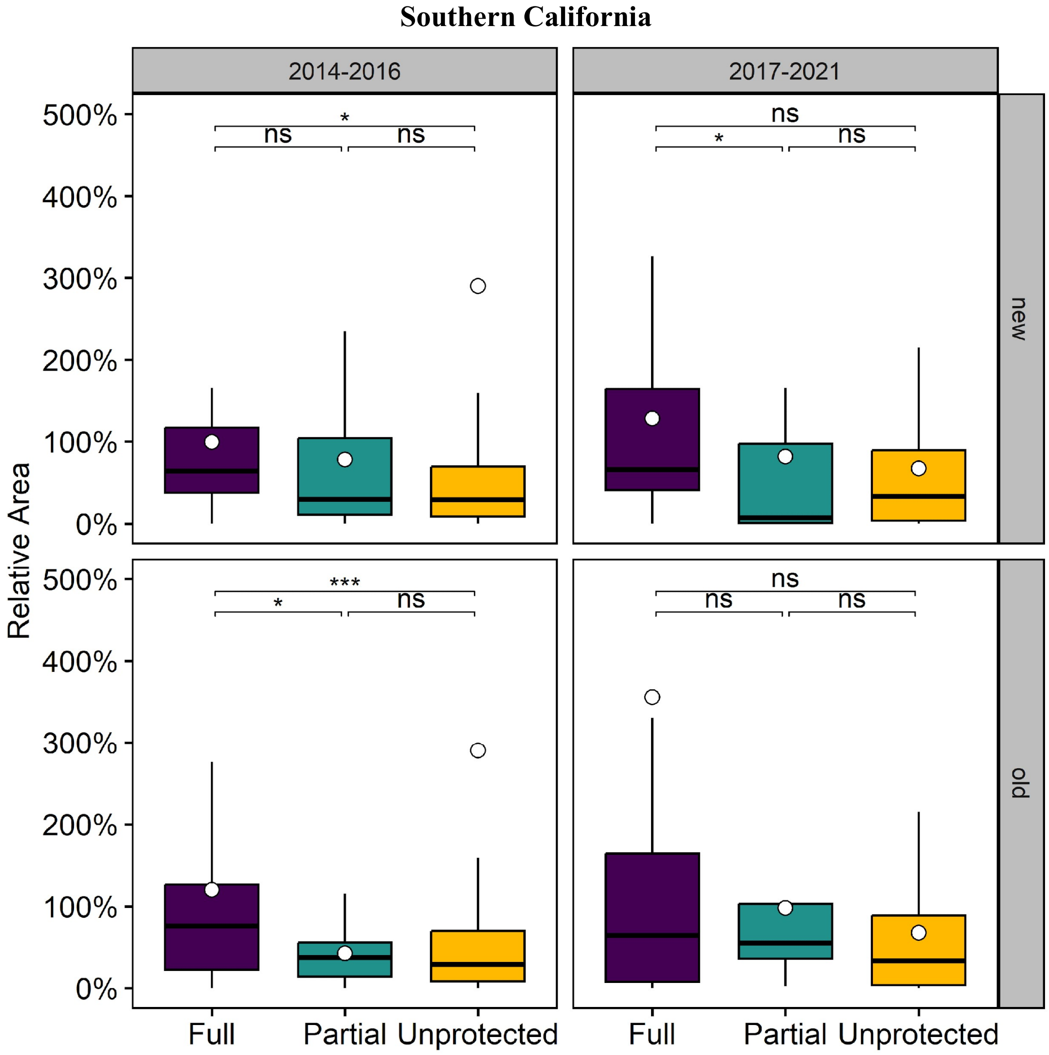


**Supplementary Figure 5**: Mean kelp canopy area relative to the historic baseline for each pixel during (2014–2016) and after the MHWs (2017–2021) in Southern California within fully protected, partially protected, and unprotected areas considering only MPAs established after 2007 (top row), and MPAs established before and during 2007 (bottom row). White points represent averages per group. Outliers are also removed from the plot for ease of visualization. Pseudo p-values were computed via Bonferroni-corrected permutation analyses; non-significant group differences are indicated with "ns" while significant comparisons are denoted with asterisks – $p < 0.05$ (*), $< 0.01$ (**), and $< 0.001$ (***).

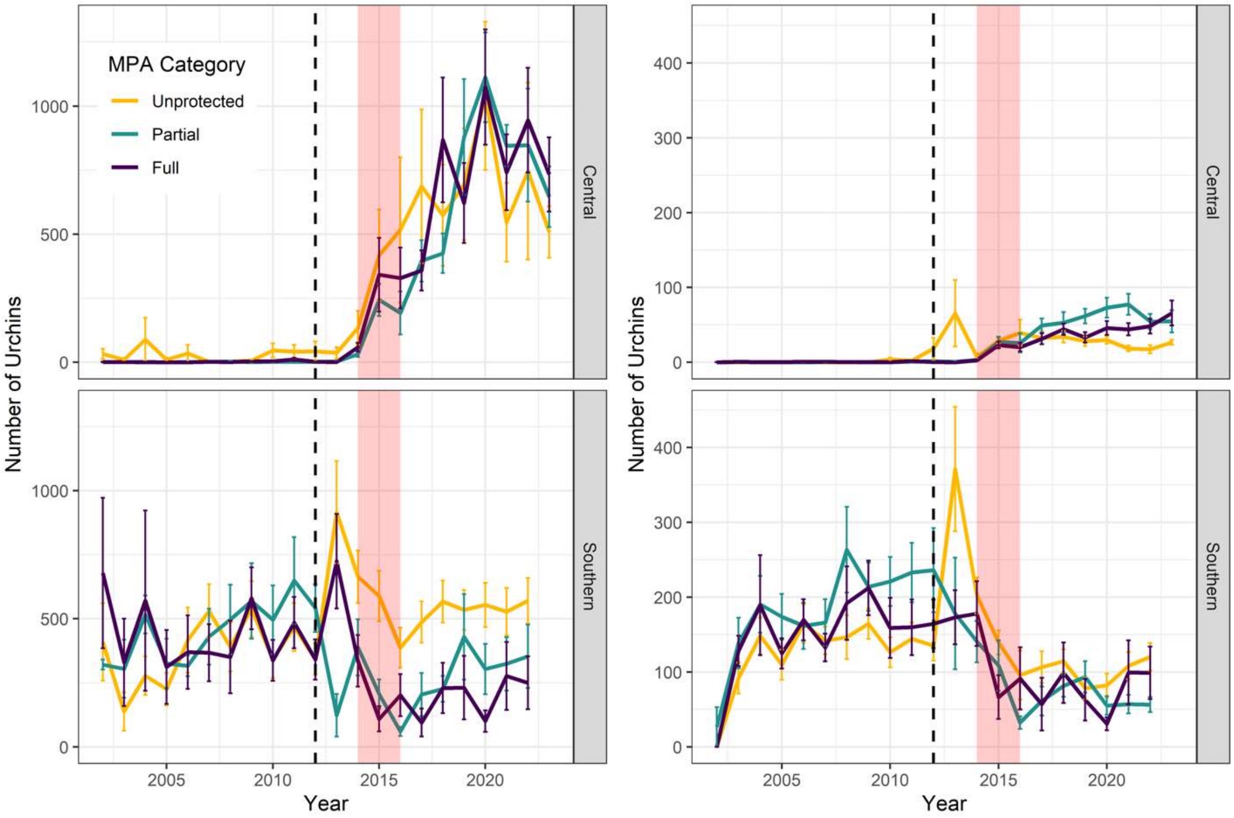


**Supplementary Figure 5:** Purple (left) and red (right) urchin abundances per standard 60m$^2$ transect through time for both Central (top) and Southern California (bottom) displayed separately. The dashed line at 2012 represents the implementation of the last marine protected areas under the Marine Life Protection Act. Data before 2012 includes sites that were protected at that time or will be protected in 2012. Data within fully protected areas are in purple, partially protected areas in turquoise, and unprotected areas in yellow. The heatwave is depicted in transparent red. Note that the scales between purple and red urchin plots are different.

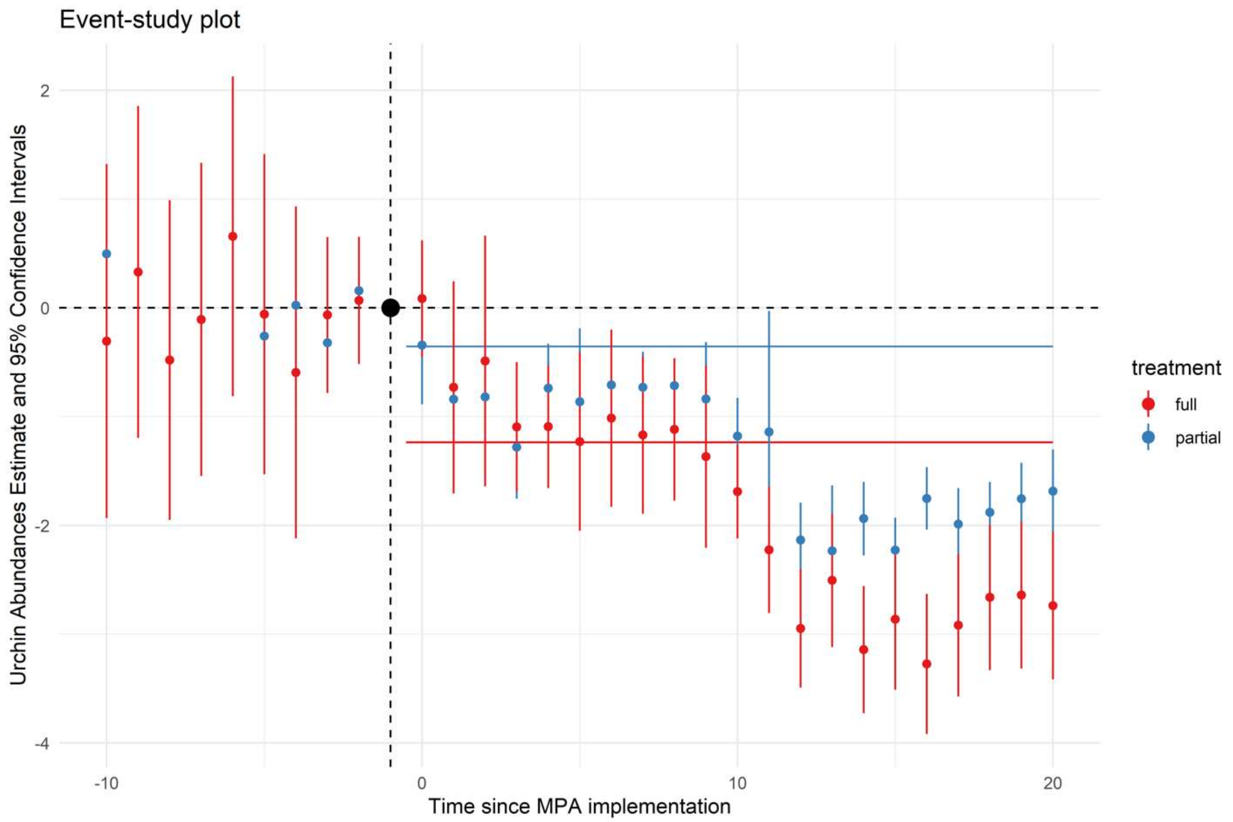


**Supplementary Figure 6:** Two-way effects model showcasing urchin abundances change over time since MPAs implementation in Southern California. This model accounts for site-level and year-level effects. The difference in urchin abundances before MPA implementation are not different from 0. The differences from the treatment increase through time, with urchin abundances decreasing as time since MPA implementation increases. The average treatment effect from the second regression is shown as solid lines and is the difference in difference estimate. These lines are the mean difference in the post-period relative to the reference level after accounting for temporal trends observed across all treatments.

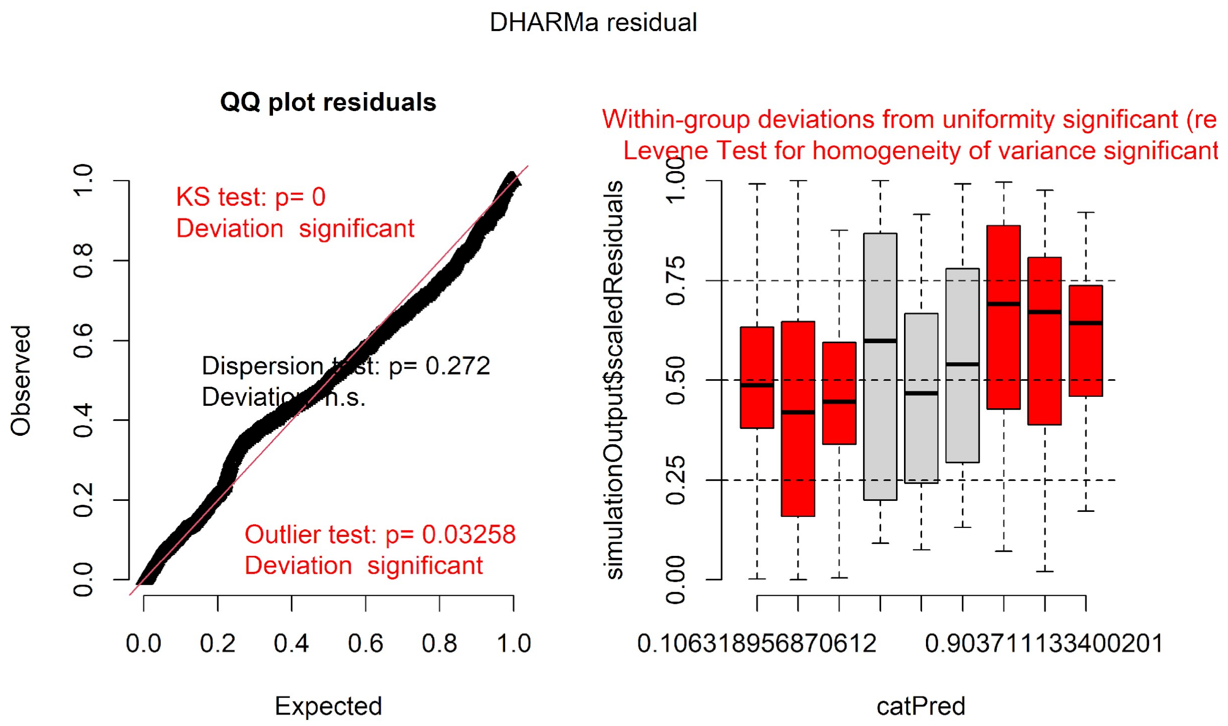


**Supplementary Figure 7:** Residuals plots from the DHARMa package for the tweedie GLMM which predicts urchin abundances based on the interaction between protection level and heatwave period in Central California (Figure 4 in the main text).

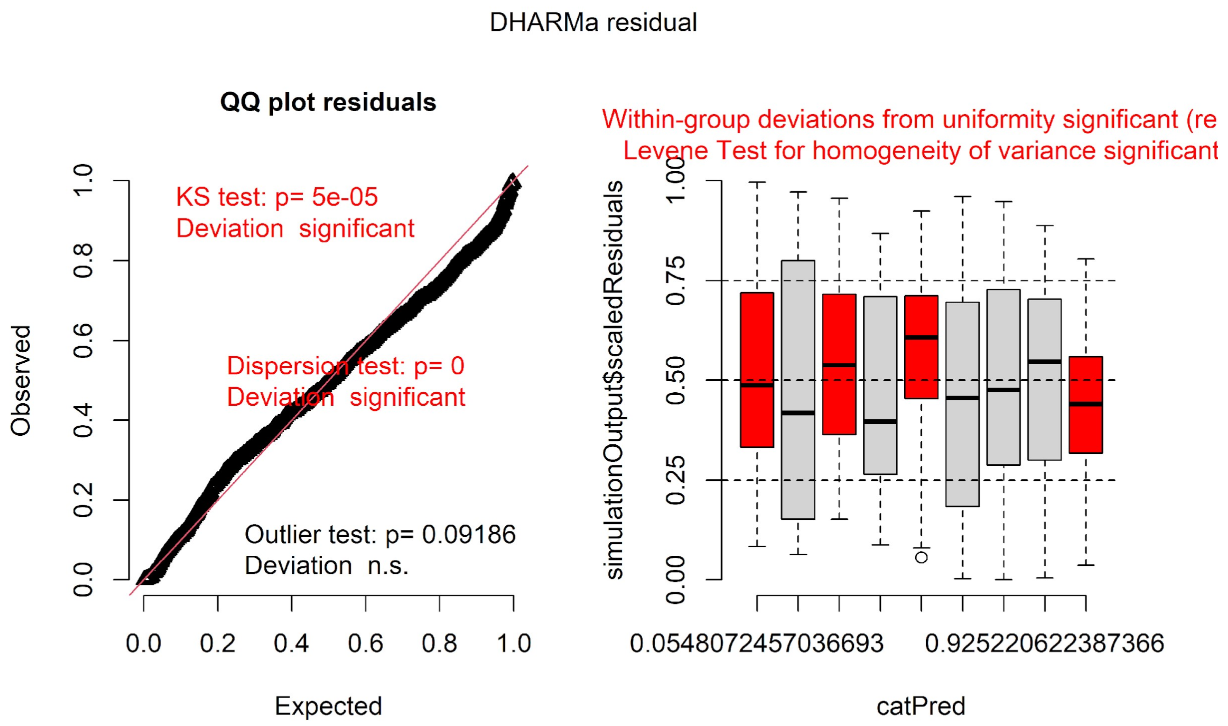


**Supplementary Figure 8:** Residuals plots from the DHARMa package for the tweedie GLMM which predicts urchin abundances based on the interaction between protection level and heatwave period in Southern California (Figure 4 in the main text).

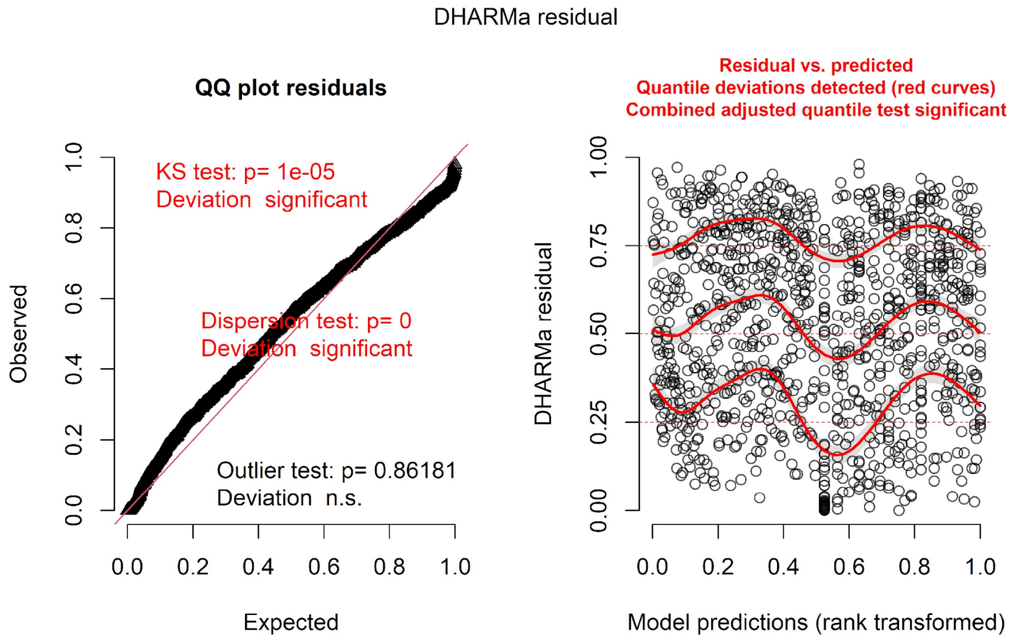


**Supplementary Figure 9:** Residuals plots from the DHARMa package for the first tweedie model which predicts urchin abundances as a product of abundances of California sheephead and spiny lobsters without an autoregressive function specified (Figure 6 in the main text, top panel).

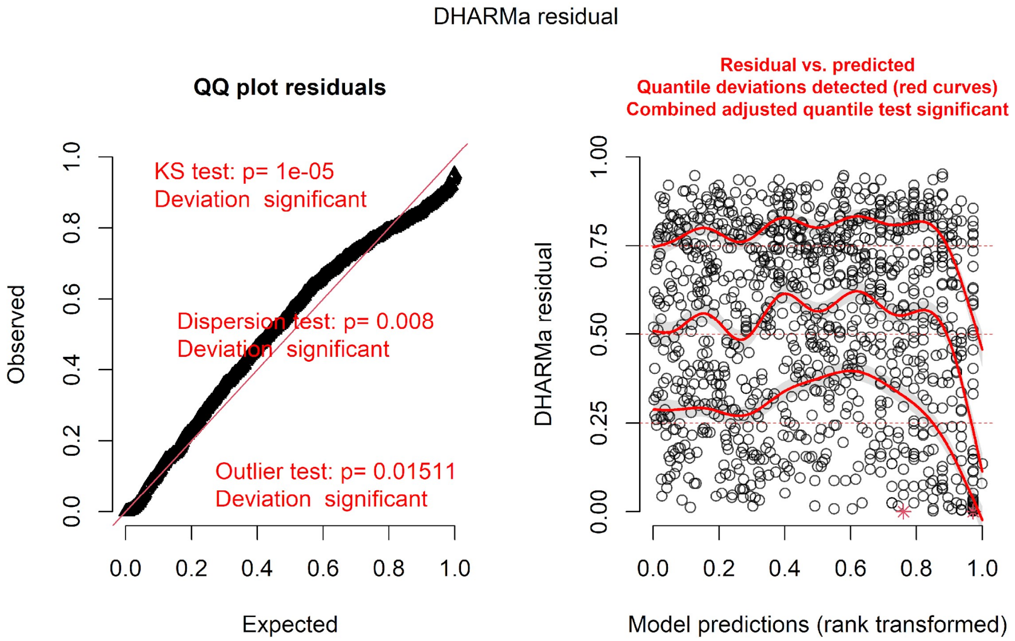

**Supplementary Figure 10:** Residuals plots from the DHARMa package for the second tweedie model which predicts urchin abundances as a product of abundances of California sheephead and spiny lobsters with an autoregressive function specified (Figure 6 in the main text, bottom panel).

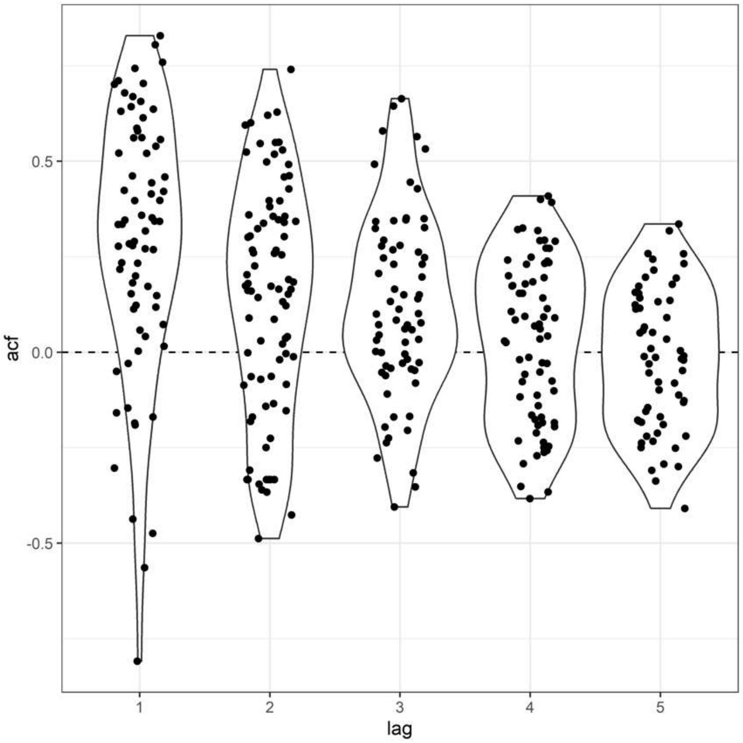


**Supplementary Figure 11:** Violin plot of the autocorrelation within the first tweedie model of urchin density predicted by California sheephead and spiny lobster without the autoregressive function specified. There was overall very little autocorrelation within the first model (Figure 6, top panel).

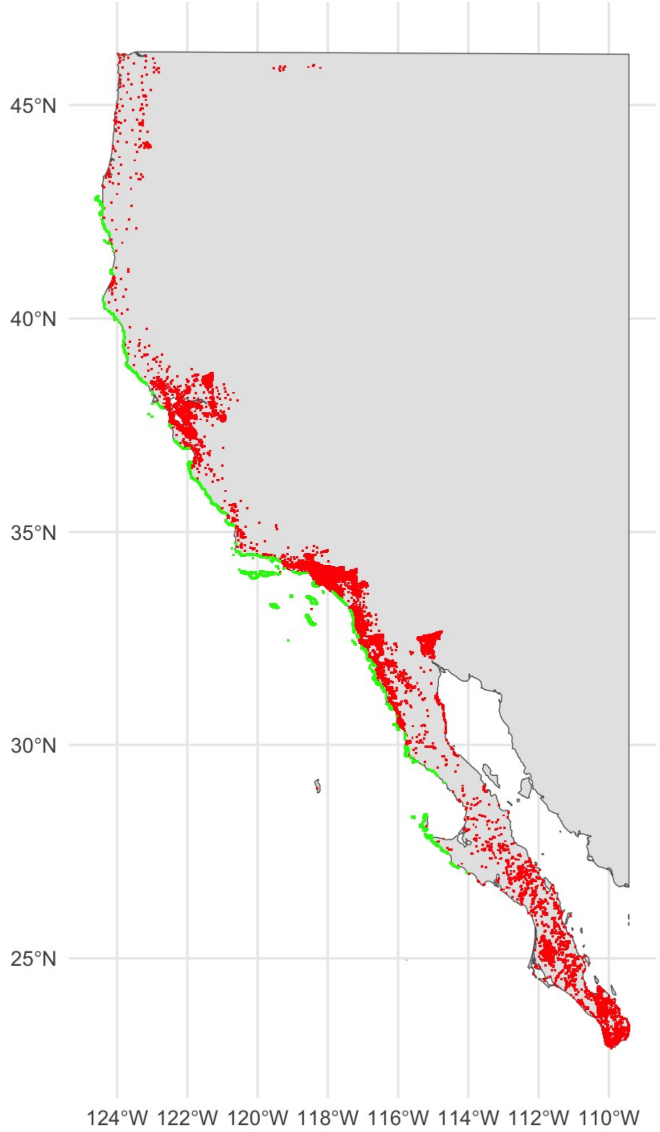


**Supplementary Figure 12:** Kelp and human settlements. Green points show places of known kelp presence. Red points are human population settlements that are within 50km of the coast.

### 3) Tables

**Supplementary Table 1)** Description of each variable considered within the study. All variables were transformed to 1-km grid resolution on a yearly scale.

| **Variable** | **Description (per 1km$^2$ pixel)** | **Data Source** |
|---|---|---|
| **Kelp area** | Mean kelp area per year | Bell, T. W., 2023 |
| **Surface Nitrate** | Mean modeled nitrate at the sea surface | Bell, T. W., 2023 |
| **Maximum significant wave height** | Mean modeled maximum wave height per year | Bell, T. W., 2023 |
| **Temperature** | Average sea surface temperature | Bell, T. W., 2023 |
| **Cumulative marine heat wave intensity** | Cumulative degrees above the 90$^{th}$ percentile of sea surface temperature during MHW events | Derived from NOAA's 0.25 degree OISST |
| **Cumulative marine cold spell intensity** | Cumulative degrees below the 10$^{th}$ percentile of sea surface temperature during marine cold-spell events | Derived from NOAA's 0.25 degree OISST |
| **Marine protected area status** | Category of either not protected ("None"), within a protected that allows some extractive activities ("Partial"), or protected with no extractive activities allowed ("Full") | Derived from NOAA's MPA Inventory |
| **Depth** | Mean depth of the pixel | Bell, T. W., 2023 |
| **Human Gravity Index** | Density-weighted exposure of each kelp patch to human population, and measured in # of people / km$^2$ | Derived from our kelp data layer, and human population from the 2020 Mexican and US Census. |

**Supplementary Table 2) Results from the permutation analysis comparing differences in relative changes in kelp area during and after the 2014–2016 MHWs between all combinations of protection categories.** Each value within the table is a pseudo *p*-value from the analysis and is adjusted for the six comparisons within each region.

| **Comparison** | **p-value 2014–2016** | **p-value 2017–2021** |
|---|---|---|
| *Central and Southern California* | | |
| Full vs Unprotected | 0 | 0.0024 |
| Partial vs Unprotected | 0.259 | 1 |
| Full vs Partial | 0.407 | 0.417 |
| *Central California* | | |
| Full vs Unprotected | 1 | 1 |
| Partial vs Unprotected | 0.02 | 1 |
| Full vs Partial | 1 | 1 |
| *Southern California* | | |
| Full vs Unprotected | 0.0006 | 0.027 |
| Partial vs Unprotected | 1 | 1 |
| Full vs Partial | 0.0054 | 0.0348 |

**Supplementary Table 3) Estimated marginal means for urchin abundances derived from the generalized linear mixed model used to predict total urchins using the heatwave period and protection category in each region.** These values are back transformed on the log scale.

| **Southern California** | | | |
|---|---|---|---|
| *Heatwave Period* | *Protection Category* | *Response (N of urchins)* | *Standard Error* |
| Before | Unprotected | 418 | 73 |
| Before | Partial | 729 | 246 |
| Before | Full | 419 | 93 |
| During | Unprotected | 587 | 107 |
| During | Partial | 302 | 103 |
| During | Full | 182 | 43 |
| After | Unprotected | 418 | 79 |

| | | | |
|---|---|---|---|
| After | Partial | 228 | 78 |
| After | Full | 83 | 21 |
| **Central California** | | | |
| *Heatwave Period* | *Protection Category* | *Response (N of urchins)* | *Standard Error* |
| Before | Unprotected | 3 | 1 |
| Before | Partial | 3 | 1 |
| Before | Full | 2 | 1 |
| During | Unprotected | 99 | 32 |
| During | Partial | 136 | 49 |
| During | Full | 139 | 41 |
| After | Unprotected | 237 | 84 |
| After | Partial | 519 | 176 |
| After | Full | 414 | 119 |

**Supplementary Table 4) Pairwise comparisons for the generalized linear mixed model used to predict total urchins using the heatwave period and protection category in each region.** *P*-values are adjusted using the tukey method for comparing 3 estimates. These tests are performed on the log scale. *P*-values lower than 0.05 are in red.

| **Southern California** | | | | |
|---|---|---|---|---|
| *Heatwave Period* | *Comparison* | *Ratio* | *Standard Error* | *P-value* |
| Before | Unprotected / Partial | 0.57 | 0.22 | 0.300 |
| Before | Unprotected / Full | 1.00 | 0.27 | 1.000 |
| Before | Partial / Full | 1.74 | 0.69 | 0.339 |
| During | Unprotected / Partial | 1.95 | 0.74 | 0.188 |
| During | Unprotected / Full | 3.23 | 0.93 | 0.000 |
| During | Partial / Full | 1.66 | 0.68 | 0.426 |
| After | Unprotected / Partial | 1.83 | 0.71 | 0.259 |
| After | Unprotected / Full | 5.06 | 1.52 | 0.000 |

| | | | | |
|---|---|---|---|---|
| After | Partial / Full | 2.76 | 1.14 | 0.039 |
| **Central California** | | | | |
| *Heatwave Period* | *Comparison* | *Ratio* | *Standard Error* | *P-value* |
| Before | Unprotected / Partial | 0.91 | 0.42 | 0.980 |
| Before | Unprotected / Full | 1.07 | 0.43 | 0.983 |
| Before | Partial / Full | 1.17 | 0.58 | 0.942 |
| During | Unprotected / Partial | 0.72 | 0.31 | 0.731 |
| During | Unprotected / Full | 0.71 | 0.27 | 0.641 |
| During | Partial / Full | 0.98 | 0.41 | 0.998 |
| After | Unprotected / Partial | 0.46 | 0.18 | 0.127 |
| After | Unprotected / Full | 0.57 | 0.21 | 0.283 |
| After | Partial / Full | 1.25 | 0.49 | 0.830 |

**Supplementary Table 5) Mean and median relative area of kelp (averaged yearly kelp area divided by the historic baseline area within each pixel) are reported within each combined category of region, heatwave period, and protection category.** Full boxplots are shown in figure 2 in the main text**.**

| **Central and Southern California** | | | |
|---|---|---|---|
| *Heatwave Period* | *Protection Category* | *Median* | *Mean* |
| During | Unprotected | 0.405 | 2.76 |
| During | Partial | 0.589 | 2.05 |
| During | Full | 0.742 | 2.14 |
| After | Unprotected | 0.479 | 3.02 |
| After | Partial | 0.551 | 4.62 |
| After | Full | 0.757 | 4.77 |
| **Central California** | | | |
| *Heatwave Period* | *Protection Category* | *Median* | *Mean* |
| During | Unprotected | 0.703 | 2.52 |
| During | Partial | 1.27 | 3.58 |
| During | Full | 0.742 | 3.17 |
| After | Unprotected | 0.731 | 7.32 |
| After | Partial | 0.789 | 8.93 |
| After | Full | 0.813 | 6.90 |
| **Southern California** | | | |
| *Heatwave Period* | *Protection Category* | *Median* | *Mean* |
| During | Unprotected | 0.290 | 2.90 |
| During | Partial | 0.295 | 0.710 |
| During | Full | 0.646 | 1.12 |
| After | Unprotected | 0.331 | 0.672 |
| After | Partial | 0.231 | 0.851 |
| After | Full | 0.645 | 2.67 |

**Supplementary Table 6) Adjusted pseudo p-values for the sensitivity analysis of differences in percent recovery between protection categories.** We investigated how sensitive the analysis presented in figure 2 was to very high percent recovery values due to low mean kelp area during 1984 to 2013. We removed pixels where the historic kelp area was lower than the 5%, 10%, 15%, 20%, 25%, and 30% quantile of all of the data from 1984 to 2013 and re-ran the permutation analysis for both regions combined and separetly.

| Quantile | Comparison | Both Regions | | South | | Central | |
|---|---|---|---|---|---|---|---|
| | | 2014-2016 | 2017-2021 | 2014-2016 | 2017-2021 | 2014-2016 | 2017-2021 |
| 5% | Full vs. None | 0.0012 | 0.0750 | 0 | 0.0588 | 1 | 1 |
| 5% | Partial vs. None | 0.2718 | 1 | 1 | 1 | 0.0246 | 1 |
| 5% | Full vs. Partial | 1 | 0.9828 | 0.006 | 0.111 | 1 | 1 |
| 10% | Full vs. None | 0.0006 | 0.1650 | 0 | 0.102 | 1 | 1 |
| 10% | Partial vs. None | 0.2070 | 1 | 1 | 1 | 0.0198 | 1 |
| 10% | Full vs. Partial | 1 | 1 | 0.0084 | 0.1218 | 1 | 1 |
| 15% | Full vs. None | 0.0000 | 0.1800 | 0 | 0.177 | 0 | 0.03 |
| 15% | Partial vs. None | 0.1236 | 1 | 1 | 1 | 0.026 | 0.3836 |
| 15% | Full vs. Partial | 1 | 1 | 0.0108 | 0.1446 | 0.1809 | 0.1832 |
| 20% | Full vs. None | 0.0000 | 0.3408 | 0.0006 | 0.1788 | 1 | 1 |
| 20% | Partial vs. None | 0.1446 | 1 | 1 | 1 | 0.0132 | 1 |
| 20% | Full vs. Partial | 1 | 1 | 0.0186 | 0.204 | 1 | 1 |
| 25% | Full vs. None | 0.0030 | 0.3516 | 0 | 0.086 | 1 | 1 |
| 25% | Partial vs. None | 0.1854 | 1 | 1 | 1 | 0.0078 | 1 |
| 25% | Full vs. Partial | 1 | 1 | 0.006 | 0.306 | 1 | 1 |
| 30% | Full vs. None | 0.0144 | 0.3762 | 0 | 0.1068 | 1 | 1 |
| 30% | Partial vs. None | 0.0024 | 0.5076 | 0.3822 | 1 | 0.0066 | 1 |
| 30% | Full vs. Partial | 1 | 1 | 0.1656 | 0.672 | 1 | 1 |
